# Effects of injury size on local and systemic immune cell dynamics in volumetric muscle loss

**DOI:** 10.1101/2024.08.26.609702

**Authors:** Ricardo Whitaker, Samuel Sung, Tina Tylek, Gregory Risser, Erin O’Brien, Phoebe Ellin Chua, Thomas Li, Ryan J. Petrie, Lin Han, Benjamin Binder-Markey, Kara L. Spiller

## Abstract

We took a systems approach to the analysis of macrophage phenotype in regenerative and fibrotic volumetric muscle loss outcomes in mice together with analysis of systemic inflammation and of other leukocytes in the muscle, spleen, and bone marrow. Macrophage dysfunction in the fibrotic group occurred as early as day 1, persisted to at least day 28, and was associated with increased numbers of leukocytes in the muscle and bone marrow, increased pro-inflammatory marker expression in splenic macrophages, and changes in the levels of pro-inflammatory cytokines in the blood. The most prominent differences were in muscle neutrophils, which were much more abundant in fibrotic outcomes compared to regenerative outcomes at day 1 after injury. However, neutrophil depletion had little to no effect on macrophage phenotype or on muscle repair outcomes. Together, these results suggest that the entire system of immune cell interactions must be considered to improve muscle repair outcomes.

## 1. Introduction

Volumetric Muscle Loss (VML) is a debilitating condition resulting from the loss of a large portion of muscle due to injury or disease. The lifetime projected disability-related expenses is greater than $400,000 USD per patient ^1^. The current standard of care for VML patients is physical therapy, which has limited benefits ^2,3^. While xenogeneic decellularized extracellular matrix (ECM) scaffolds have shown promise in clinical trials, the results still fall far short of restoring pre-VML functionality ^4,5^. There is a major need for more effective interventions that can promote regenerative tissue healing.

Recent studies have sought to identify a cellular basis for dysfunctional wound healing following VML. Several studies have pointed to macrophages, which are major regulators of wound healing and primary targets to improve tissue regeneration across a wide variety of tissues (reviewed in Ref. ^6,7^). A series of landmark studies showed that implantation of ECM-derived scaffolds promoted regenerative outcomes of VML in mice, associated with skewing of macrophages towards a Th2-associated profile ^8–10^. On the other hand, macrophages in fibrotic outcomes generated by poly(caprolactone) (PCL) implants were skewed towards a Th17 profile. Other important studies used critical sized defects in the absence of biomaterial implantation to generate fibrotic outcomes in comparison to smaller injuries that almost perfectly regenerate ^11–16^. In general, these studies showed that macrophages in fibrotic injuries simultaneously expressed high levels of pro-inflammatory factors like tumor necrosis factor-alpha (TNFα) and pro-fibrotic factors like transforming growth factor-beta (TGFβ) ^11,15^. At the same time, macrophages in fibrotic injuries expressed lower levels of markers previously associated with regenerative macrophage behavior in other tissues. These markers included CD206, CD301b, CD163, and PDGFBB ^10,15,16^, although at least one study noted elevated levels of CD206+ macrophages in fibrotic VML ^11^. However, it is not known for how long these phenotype changes persist, and the mechanisms behind them are not known, making it difficult to design targeted interventions that correct macrophage behavior for improved tissue regeneration.

The behavior of other infiltrating immune cells to injury sites can influence macrophage phenotype. For example, Sadtler et al. showed that the phenotype of infiltrating T cells are critical for promotion of a regenerative macrophage phenotype in ECM scaffold-promoted regeneration of VML ^17^. However, it is not clear to what extent this effect was specific to the ECM scaffold microenvironment or whether it is true for regenerative outcomes in the absence of ECM scaffolds. Neutrophils have been shown in several disease and injury models to directly influence macrophage phenotype through efferocytosis, which promotes an anti-inflammatory and regenerative macrophage phenotype ^18–22^, and secretion of soluble factors (which can promote a pro-inflammatory macrophage phenotype) ^23–25^. Although the effects of neutrophils on macrophage phenotype in VML have not been thoroughly investigated, one study did show that the prolonged presence of neutrophils was detrimental for muscle repair via effects on satellite cells ^14^. To our knowledge, no study has employed a holistic approach to analyze macrophage phenotype in the context of the many immune cell interactions within this system. It is critical to understand how macrophage phenotype dynamics change with other immune cells to design targeted interventions to enhance tissue repair.

Finally, it is not clear to what extent VML influences the entire immune system outside of the local muscle tissue itself. In major traumatic injury in humans, systemic levels of pro-inflammatory cytokines are known to be elevated in the bloodstream ^26–29^. In mice, systemic effects of injuries including myocardial infarction, stroke, and sepsis have been shown to influence the behavior of macrophages in other organs like heart and brain ^30^. In that study, the behavior of immune cells in the reservoir organs (i.e. bone marrow and spleen) was not evaluated. Spleen- and bone marrow-derived immune cells are the principal sources of monocyte-derived macrophages and other leukocytes that populate sites of injury ^31,32^. These studies suggest that critical size muscle defects might have systemic influence on immune cells at the reservoir organs, with possible repercussions on leukocyte recruitment to the site of injury. Therefore, the purpose of this study was to evaluate the local and systemic changes in immune cell behavior in fibrotic vs. regenerative muscle repair, with the goal of identifying possible mechanisms of macrophage dysfunction in fibrotic outcomes. We hypothesized that the magnitude of the injury in fibrotic muscle repair has systemic repercussions that result in dysfunctional behavior in monocyte-derived macrophages, ultimately leading to worse healing outcomes.

To investigate how the size of the injury influences local and systemic changes in immune cell behavior, we utilized a murine model of VML in which small (2mm) defects in the quadriceps completely regenerate while critically sized (4mm) defects result in fibrotic repair with both reduced quantity and quality of repair tissue ^33^. We characterized local macrophage phenotype and other immune cell profiles in the muscle via multi-dimensional flow cytometry and multiplex gene expression analysis (nanoString). Then, we investigated systemic effects of the two injury types on immune cell behavior and macrophage phenotype at the major immune cell reservoirs, the bone marrow and the spleen, using flow cytometry, and we measured systemic levels of immunomodulatory factors in blood serum through multiplex cytokine analysis (Luminex). Finally, we investigated whether reducing the levels of neutrophils to those observed in the smaller injury size would influence macrophage phenotype. These findings provide critical insight into the mechanisms behind dysfunctional immune cell dynamics in fibrotic muscle injuries.

## 2. Results

### 2.1. Dysfunctional macrophage behavior in fibrotic muscle repair

To evaluate the mechanism driving pathological macrophage phenotype following VML, we utilized a murine model in which a critically sized defect (4mm) in the quadriceps leads to fibrotic outcomes, while smaller defects (2mm) lead to regeneration ^14,33^ (**Figure 1A, Suppl. Figure 1**). By 28 days, the muscle tissue in the fibrotic group was characterized by higher levels of collagen staining, sulfated glycosaminoglycans (sGAGs) content, and overall muscle mass compared to the regenerative group and to naïve (uninjured) tissue, while no differences were observed between the regenerative and the naïve groups (**Figure 1B**). We next characterized the local and systemic immune response in these groups via multidimensional flow cytometry, multiplex gene expression analysis (nanoString) (**Suppl. Table 1**), multiplex cytokine analysis (Luminex), and histology and immunohistochemistry over 28 days post injury (**Figure 1A**).

**Figure 1.**
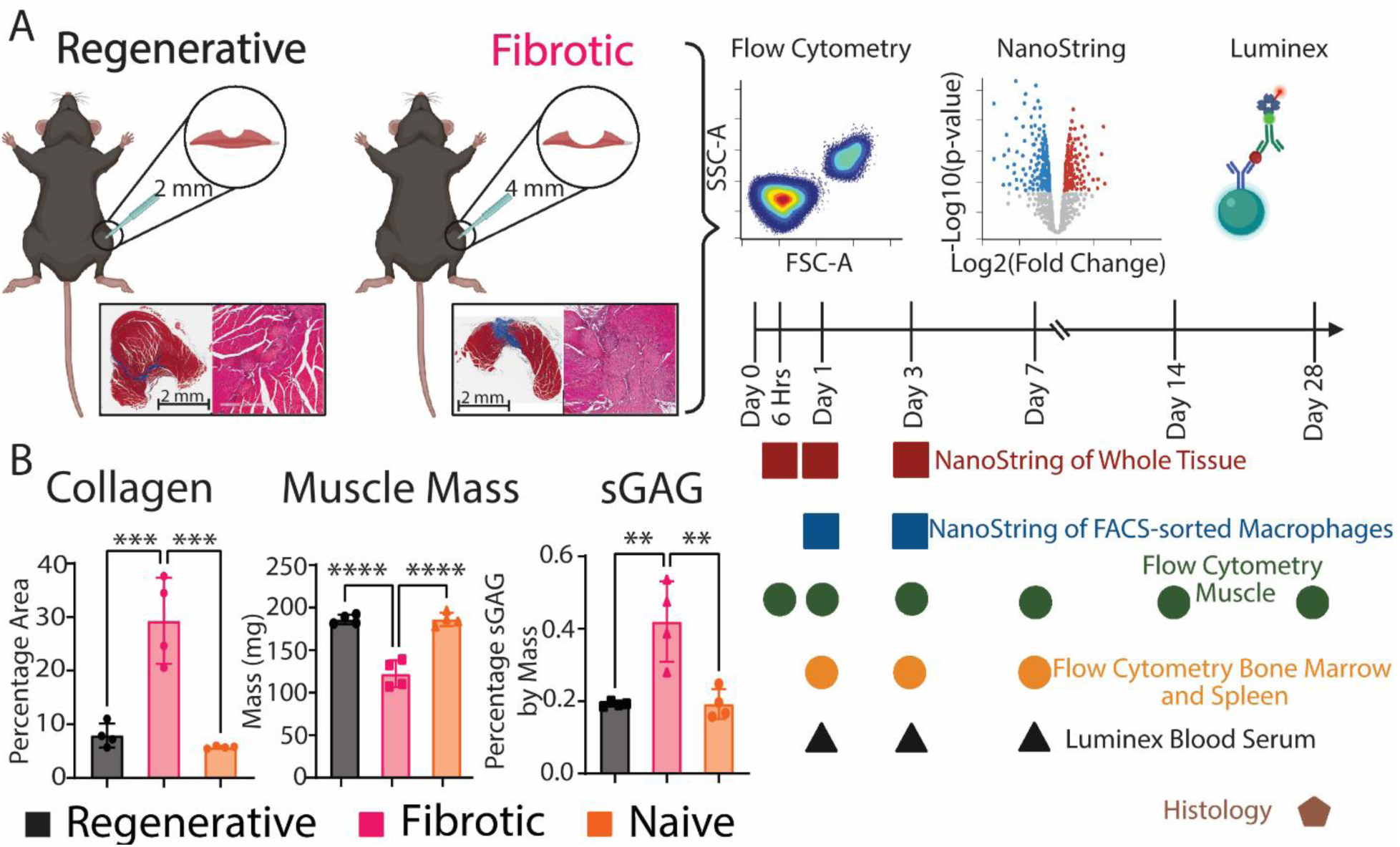
Experimental design. **A)** Overview of the model and representative images of Masson’s trichrome staining. Image created in BioRender.com. **B)** Area of collagen per section (Masson’s trichrome), overall muscle mass, and sulfated glycosaminoglycans (sGAGs) content. One way ANOVA, Tukey’s post hoc. N = 6, mean ± SD. * p<0.0.5; ** p < 0.01; *** p < 0.001; **** p < 0.0001.

The numbers of total myeloid cells (CD45+) and macrophages (CD45+B220-CD3-NK1.1-Ly6G-F4/80+) in the muscle increased dramatically after injury in both the regenerative and fibrotic outcome groups, with higher numbers in the fibrotic group at days 3 and 28 (**Figure 2A-C)**. Interestingly, as a percentage of CD45+ cells, there were slightly fewer macrophages in the fibrotic group at day 1 compared to the regenerative group (**Figure 2C**). The phenotypes of macrophages in the muscle were characterized via flow cytometry using a panel of 9 common macrophage phenotype markers, which we found to be differentially regulated by macrophages treated in vitro with pro-inflammatory (Th1), Th2, or Th17 stimuli (**Suppl. Figs. 2-4**). While overall temporal trends in marker expression were similar between groups, the magnitude of marker expression levels differed in macrophages from the regenerative or the fibrotic outcome groups as early as day 1 and persisted until day 28 (**Figure 2D, Suppl. Figure 3**).

**Figure 2.**
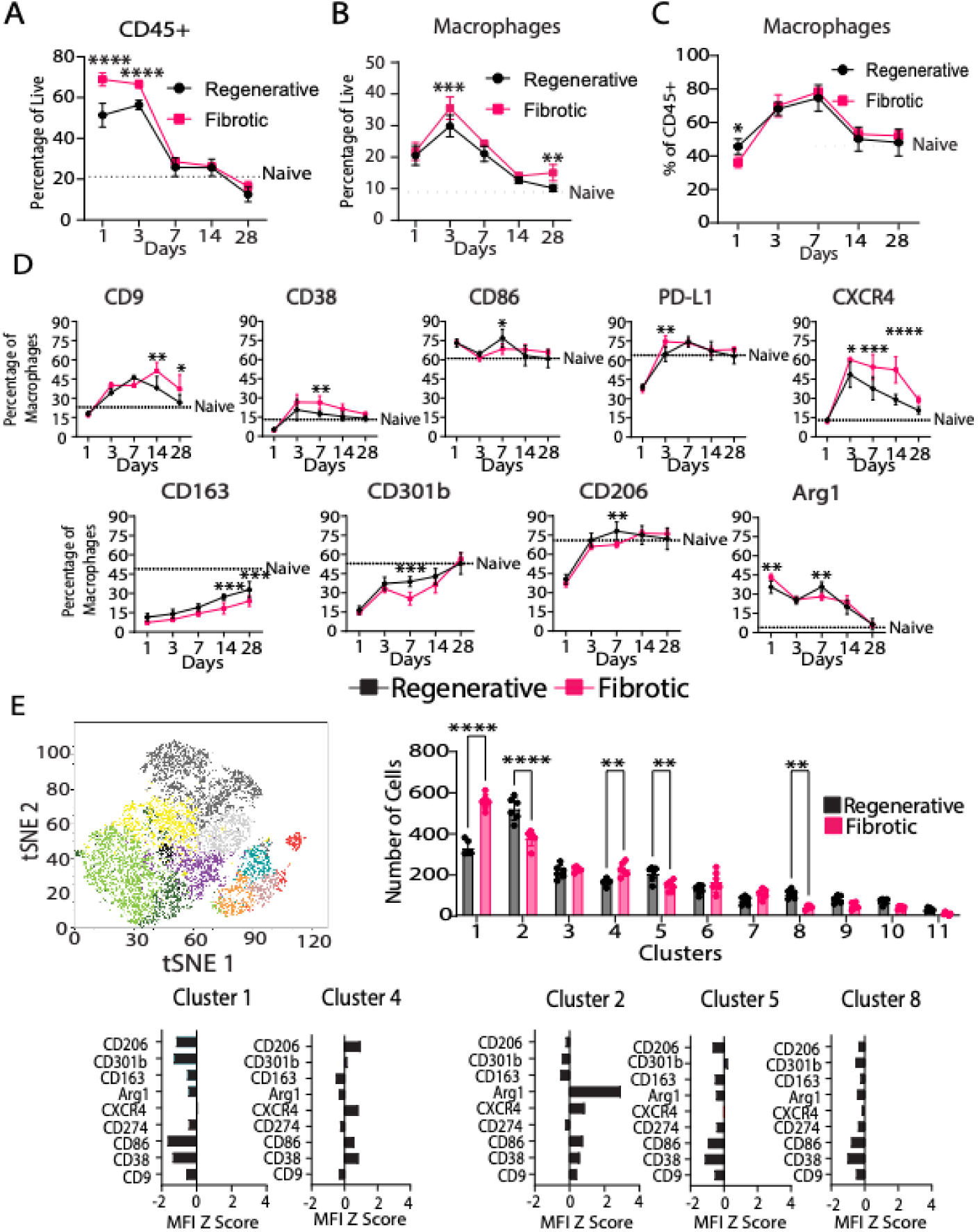
Expression levels of macrophage phenotype markers over 28 days differ between the regenerative and fibrotic outcome groups. **A)** CD45+ cells presence in muscle. **B)** Macrophage presence in muscle as a function of all cells. **C)** Macrophage presence in muscle as a function of CD45+ cells. **D)** Muscle macrophage phenotype measured by flow cytometry. **E)** Hierarchical clustering of MFI values at day 7. Two-way ANOVA, Sidak’s post hoc test. N = 6, bars show mean ± SD. * p<0.0.5; ** p < 0.01; *** p < 0.001; **** p < 0.0001.

Compared to macrophages in the regenerative group, macrophages in the fibrotic outcome group generally expressed higher levels of CD9 (previously associated with fibrotic macrophages), CD38 and PDL1 (pro-inflammatory markers that have anti-inflammatory activity), Arg1 (reparative marker involved in collagen production), and CXCR4 (involved in cell recruitment), depending on the time point (**Figure 2D, Suppl. Figure 3**). Macrophages in the fibrotic group also expressed lower levels of CD86 (pro-inflammatory marker involved in antigen presentation), CD163, CD301b, and CD206 (regenerative markers involved in phagocytosis). Macrophages in the fibrotic group exhibited a phenotype that did not resemble any of the reference phenotypes characterized in vitro (**Suppl. Figure 4**).

To uncover macrophage subpopulations, we employed multidimensional analysis via XShift and FlowSOM clustering algorithms at days 1, 3 and 7 post injury (**Figure 2E, Suppl. Fig 5**). While few differences between groups were observed at days 1 and 3 (**Suppl. Figure 5**), several differences were apparent at day 7 (**Figure 2E**).

Macrophages from the fibrotic group were present in higher numbers in clusters 1 and 4, which were characterized by low levels of all markers (cluster 1) or intermediate expression of CD206, CXCR4, CD86, and CD38 (cluster 4) (**Figure 2E**). In contrast, macrophages from the regenerative group were present in higher numbers in clusters 2, 5 and 8. While clusters 5 and 8 were characterized by low expression of all the markers, cluster 2 was characterized by very high expression of Arg1 and intermediate expression of CXCR4 and CD86. These results show that macrophages in each group are heterogeneous, with multiple subpopulations existing simultaneously.

To characterize macrophage phenotype more thoroughly at the early time points, we FACS-sorted macrophages (CD45+B220-CD3-NK1.1-Ly6G-F4/80+) from these two injuries at days 1 and 3 post injury (**Figure 3A**). We employed multiplex gene expression analysis (nanoString) using a custom-curated panel containing 259 genes related to macrophage phenotype, immune cell recruitment and trafficking, phagocytosis, wound healing, angiogenesis, and fibrosis (**Suppl. Table 1**). Hierarchical clustering of differentially expressed genes (DEGs, using a cut-off of p<0.01) at day 1 showed a clear separation between the injury outcomes (**Figure 3B**). Noticeably, there was broad downregulation of genes in the macrophages from the fibrotic outcome group, with 32 out of the 35 DEGs expressed at lower levels compared to macrophages from the regenerative group (**Figure 3C**). These downregulated genes were mostly related to tissue repair, ECM remodeling and phagocytosis (**Suppl. Table 2**). The 3 genes expressed at higher levels by fibrotic macrophages were *Cxcl2* and *Cxcl3*, both which encode leukocyte-recruiting chemokines, and *Csf1*, involved in macrophage maturation.

**Figure 3.**
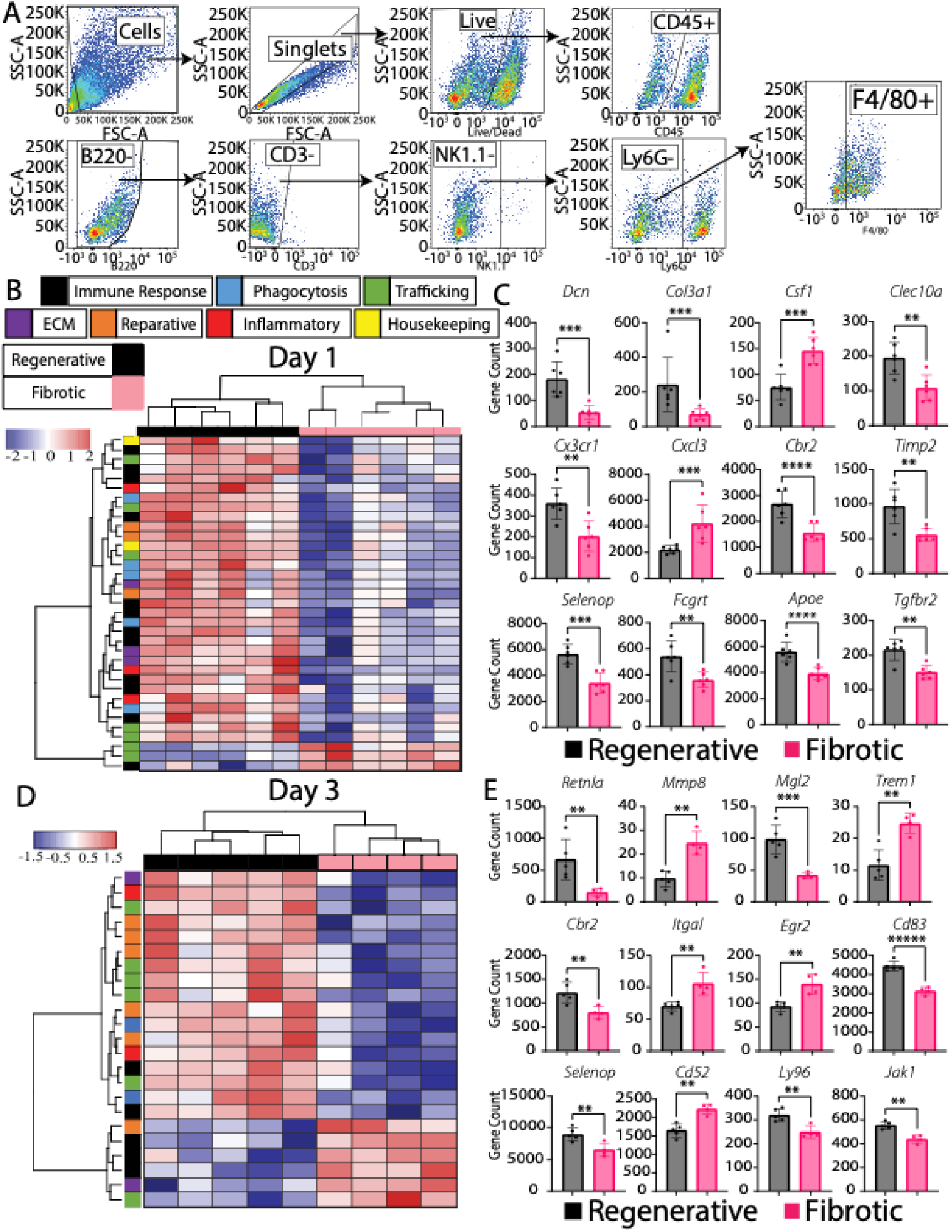
Gene expression differences in macrophage phenotype at days 1 and 3. **A)** Gating strategy for macrophage sorting. **B)** Hierarchical clustering of DEGs from nanoString multiplex gene expression analysis of FACS-sorted macrophages at day 1. **C)** Top 12 DEGs at day 1, ranked by fold change between groups. T-tests. (p<0.01 cutoff). **D)** Hierarchical clustering of DEGs from FACS-sorted macrophages at Day 3. **E)** Top 12 DEGs at day 3, ranked by fold change between groups. T-tests. (p<0.01 cutoff). N = 4 - 6, mean ± SD. ** p < 0.01; *** p < 0.001; **** p < 0.0001. Statistical analysis was performed on log-transformed data.

A similar pattern emerged at day 3, in which 17 of 23 DEGs were downregulated in the fibrotic group compared to regenerative group (**Figure 3D, Suppl. Table 2**). More than half of the downregulated genes were reparative and trafficking related genes (**Figure 3E**). The most downregulated genes in the fibrotic group were *Mgl2 (Cd301b)* and *Retnla,* which previously have been associated with regenerative behavior of macrophages in muscle and/or skin ^10,34,35^. The 6 genes that were upregulated in the fibrotic group include *Mmp8, Trem1, Itgal, Egr2, Cd52 and Ctsl*, which are involved in cell motility, metabolism, and immune responses. These results indicate that, at the gene expression level, macrophages in the fibrotic outcome group undergo major phenotype changes soon after injury compared to the regenerative outcome group.

We next characterized changes in the whole muscle tissue using the same nanoString gene expression panel. As early as 6 hours after injury, the fibrotic outcome group was characterized by higher expression of genes encoding several chemokines involved in immune cell recruitment, such as *Cxcl2* and *Cxcl3* (**Figure 4A, Suppl. Table 3).** By days 1 and 3 the fibrotic group was also characterized by increased levels of *Il36g* and *Tgfb1*, which have been previously linked to fibrotic VML ^10,13,15,16^. Finally, expression of numerous genes involved in ECM deposition and assembly were decreased in the fibrotic group at days 1 and 3, including *Col1a1*, *Col5a2*, and *Dcn*.

Given the upregulation of several chemokines associated with the trafficking of immune cells such as neutrophils and monocytes, we next evaluated immune cell populations in the muscle over 28 days (**Figure 4B, Suppl. Figure 6**). At day 1 after injury, there were nearly twice as many neutrophils in the fibrotic outcome group compared to the regenerative group, both as a percentage of leukocytes (CD45+) (**Figure 4B**) and as a percentage of all live cells (**Suppl. Figure 7**). There were also more dendritic cells in the fibrotic group at days 3, 7, and 28 when expressed as a percentage of all cells but only at day 7 when expressed as percentage of CD45+ cells. B cells, natural killer (NK) cells, monocytes and macrophages were present in lower numbers in the fibrotic group, depending on the time point (**Figure 4B, Suppl. Figure 7**).

**Figure 4.**
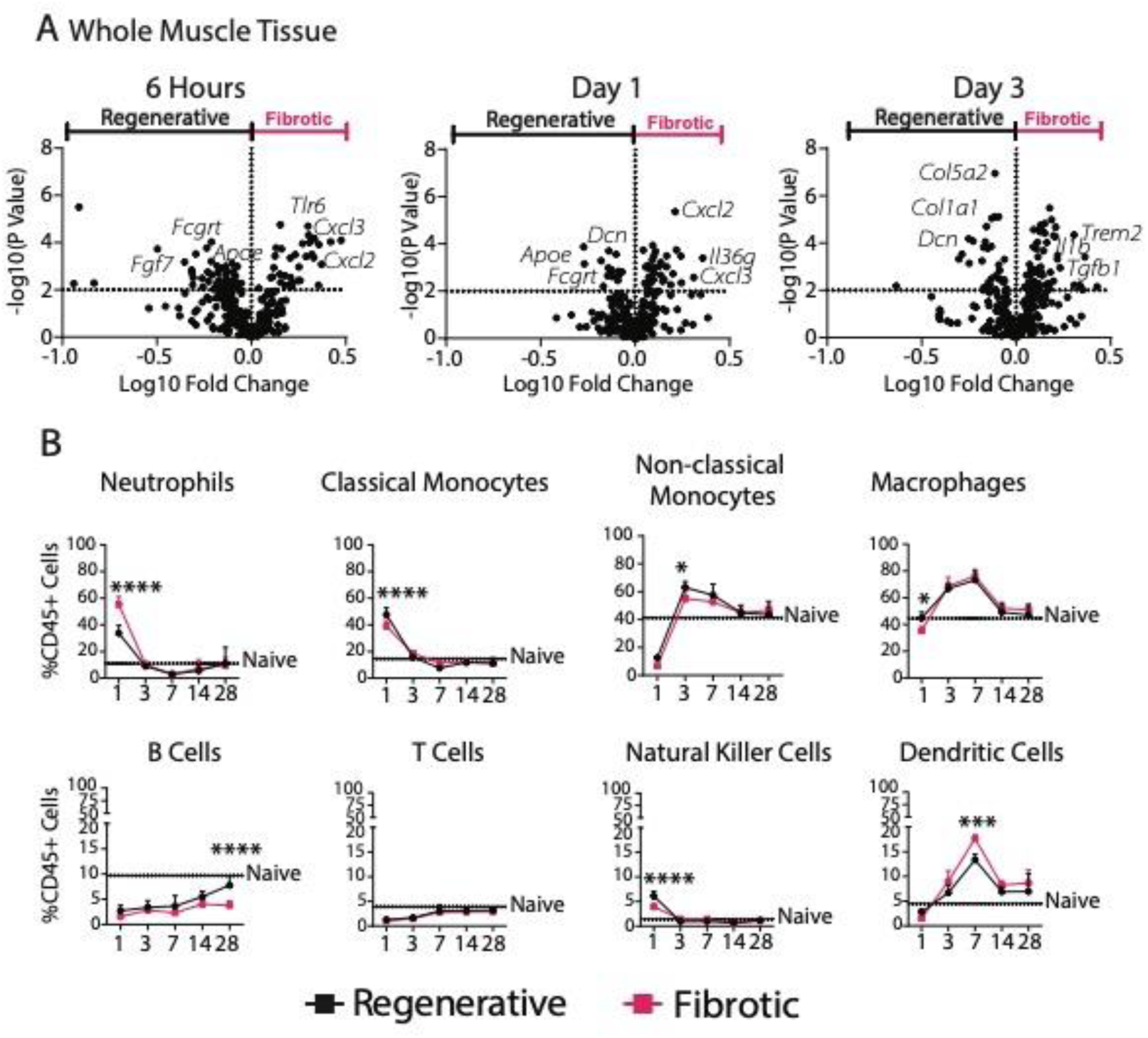
Gene expression of muscle tissue and numbers of immune cells differ in fibrotic and regenerative injuries over time. **A)** Volcano plots of nanoString from whole tissue (muscle) at 6 hours, 1 and 3 days post injury. Multiple t-tests (p<0.01 cutoff). **B)** Immune cell populations in muscle over 7 days. Two-way ANOVA, Sidak’s post hoc. N = 6, mean ± SD. * p<0.0.5; ** p < 0.01; *** p < 0.001; **** p < 0.0001.

### 2.2. Systemic changes in immune response in fibrotic vs. regenerative outcome groups

Having identified major differences in macrophage phenotype in the muscle, we next characterized the behavior of macrophages and monocytes in the spleen and bone marrow, the primary sources of monocyte-derived macrophages that infiltrate sites of injury (**Figure 5**). While there were fewer total myeloid cells (CD45+) overall in the bone marrow of mice in the fibrotic outcome group, there were more natural killer cells at day 7, and slightly more macrophages at day 3 (**Figure 5A, Suppl. Figure 8, Suppl. Figure 9**). While there were no differences in immune cell numbers in the spleen between fibrotic and regenerative outcomes (**Suppl. Figure 8**), there were several differences in macrophage phenotype in the spleen as early as day 1, persisting for up to 7 days post injury (**Figure 5B**). Spleen macrophages from the fibrotic muscle group expressed higher levels of CD38, CD86 and PDL1 and the reparative marker CD206, although these differences were relatively small. Macrophage phenotype in the bone marrow did not differ between injury types (**Suppl. Figure 8**).

**Figure 5.**
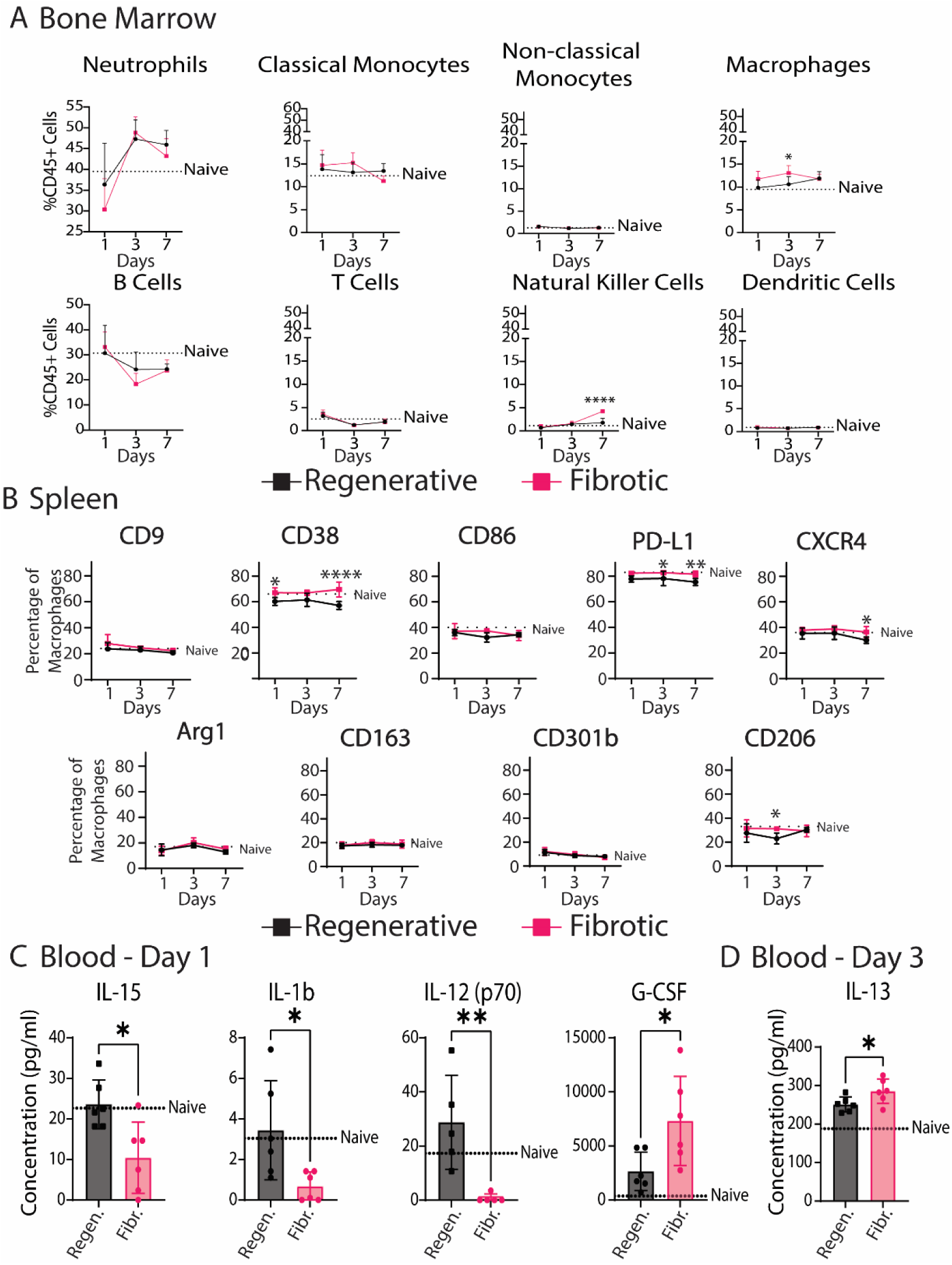
Systemic effects of injury size. **A)** Immune cell populations in bone marrow over 7 days. **B)** Macrophage phenotype in spleen over 7 days. Two-way ANOVA, Sidak’s post hoc. **C)** Significantly different analytes in blood serum at day 1. **D)** Significantly different analytes in blood serum at day 3. T-test. N = 6, bars show mean ± SD. * p<0.0.5; ** p < 0.01; *** p < 0.001; p < 0.0001 ****.

To evaluate possible signaling factors that might modulate immune cell behavior at the systemic level, we analyzed the levels of 32 immune response-related cytokines in blood serum over 7 days post injury (**Figure 5C, D, Suppl. Figure 8E, Suppl. Table 4**). At day 1, the fibrotic group was characterized by higher levels of granulocyte colony stimulating factor (G-CSF), which is crucial for neutrophil development and maintenance. There were also lower levels of pro-inflammatory cytokines interleukin-1-β (IL-1β), IL-12 (p70), and IL-15 in the fibrotic group compared to the regenerative group and to the naïve (uninjured) group. IL1β is a potently pro-inflammatory cytokine with broad effects on multiple cell types, while IL12 is involved in Th1 differentiation of T cells and IL15 is involved in natural killer (NK) cell differentiation. On day 3, the only difference was in IL-13, a fibrotic Th2 cytokine that was elevated in the fibrotic outcome group (**Figure 5D**).

### 2.3. Apoptotic neutrophils are elevated in fibrotic outcomes

Because the increased numbers of neutrophils in the fibrotic group was the greatest difference observed in our characterization thus far, we decided to further investigate their phenotype. Neutrophils have been shown to modulate macrophage phenotype via several different mechanisms, including release of soluble factors ^23^ and their phagocytosis by macrophages (efferocytosis) ^36,37^. To evaluate potential differences in neutrophil phenotype, we FACS-sorted neutrophils (CD45+B220-CD3-NK1.1-CD11b+Ly6G+) from the muscle at day 3 (**Figure 6A**). Gene expression analysis of FACS-sorted neutrophils showed 9 DEGs out of a panel of 259 between the groups (**Figure 6B**). Upregulated genes in the fibrotic group included *Pcdc1lg2*, which encodes an anti-inflammatory protein, *Lpl* and *Arg1*, which are involved in metabolism and collagen synthesis, and *Mmp13*, which is involved in ECM degradation. Upregulated genes in neutrophils from the regenerative group included *Retnla* and *Mgl2*, which have been previously associated with regenerative functions in macrophages. However, it is important to note that there was a population of Ly6G+F4/80+ cells in the muscle, so we cannot exclude possible contamination from macrophages (**Suppl. Figure 10**).

**Figure 6.**
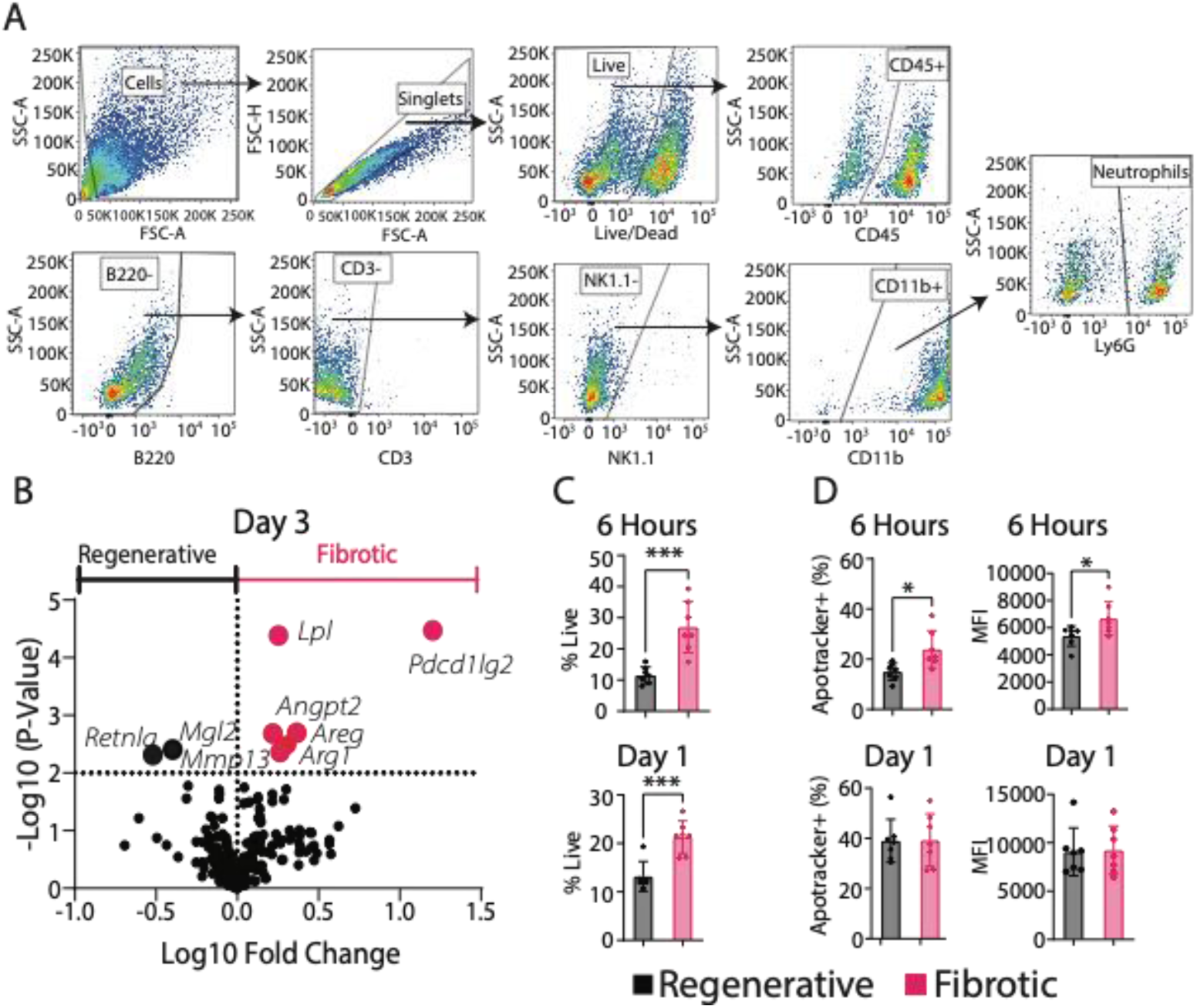
Neutrophil phenotype and levels of apoptosis. **A)** Gating strategy for neutrophil sorting. **B)** Volcano plot of nanoString gene expression data from FACS-sorted neutrophils at day 3. Multiple t-tests (p<0.01 cutoff). **C)** Neutrophil numbers in muscle at 6 hours and 1 day post injury. T-test. **D)** Neutrophil apoptosis in muscle at 6 hours and 1 day post injury as measured using ApoTracker staining in the neutrophil population. T-test. * p<0.0.5; ** p < 0.01; *** p < 0.001; **** p < 0.0001. N = 6, mean ± SD.

Next, we evaluated the levels of neutrophil apoptosis and cell death within the muscle at 6 hrs and at day 1. By 6 hours post injury, there were already increased numbers of neutrophils (**Figure 6C**) and more of them were apoptotic (**Figure 6D, Suppl. Figure 11**) in the fibrotic group compared to the regenerative group. The increased numbers of live neutrophils remained at day 1, although the percentage of neutrophils that were apoptotic did not differ between groups at this time point.

### 2.4. Partial neutrophil depletion does not affect macrophage phenotype or muscle repair outcome

Excessive levels of apoptotic neutrophils, and their impaired clearance by macrophages, have been linked to fibrosis and impaired wound healing across a range of tissues outside of VML (reviewed in Ref. ^38^). Within VML, the prolonged presence of neutrophils has been linked to persistent inflammation in fibrotic outcomes ^14^. Therefore, we hypothesized that excessive neutrophils might be a possible mediator of macrophage dysfunction and impaired tissue repair in the fibrotic outcomes. The gene expression analysis of FACS-sorted macrophages at day 3 revealed lower expression of phagocytosis- and efferocytosis-related genes in macrophages from the fibrotic group compared to the regenerative group **(Suppl. Figure 12),** suggesting possible defects in efferocytosis of neutrophils. To directly investigate the influence of excessive neutrophils on macrophage phenotype and muscle repair, we developed a partial neutrophil depletion protocol using systemic administration of anti-Ly6G antibody that depleted neutrophils in the muscle to the levels observed in the regenerative outcome group (**Figure 7A, B, Suppl. Figure 13**).

**Figure 7.**
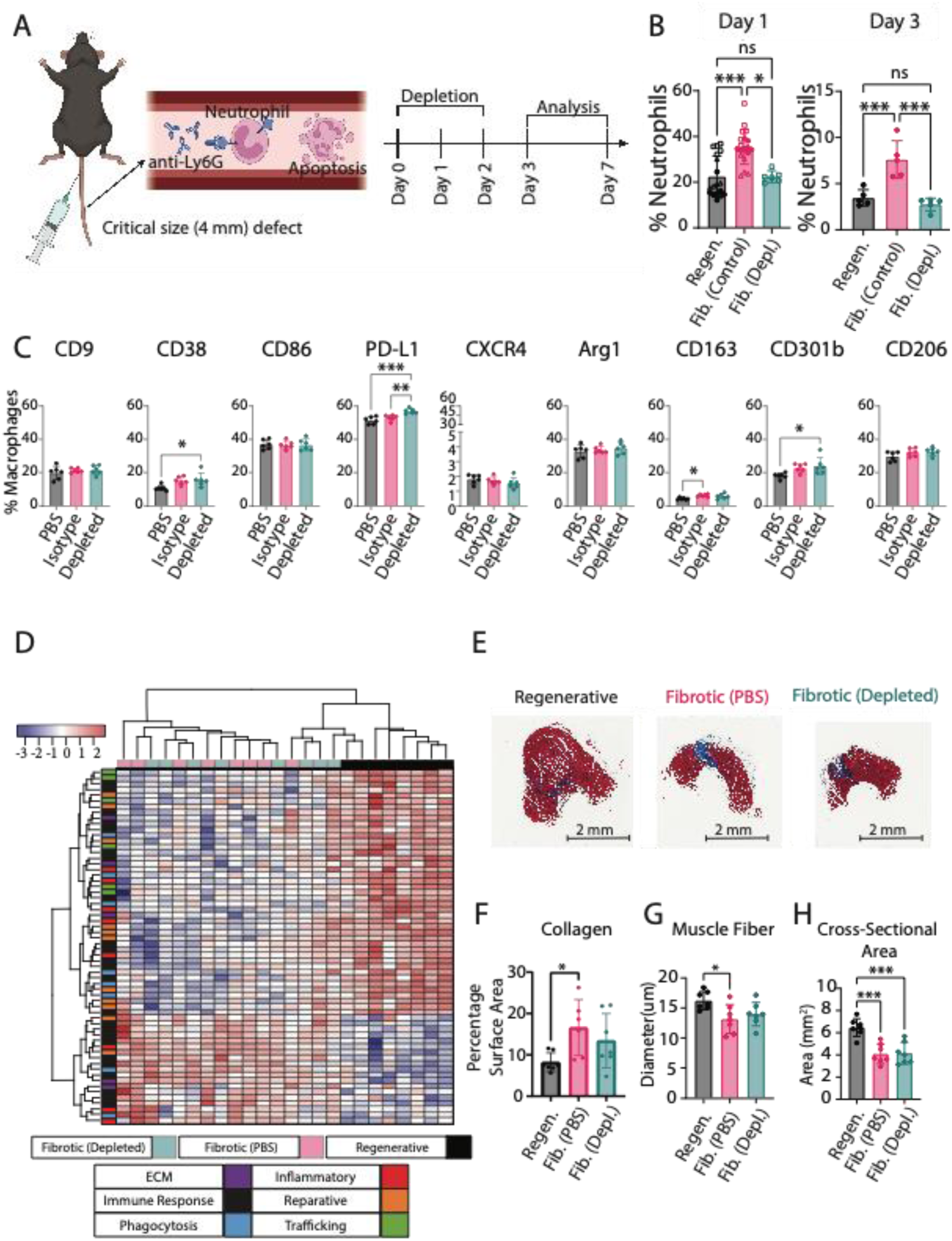
Effects of partial neutrophil depletion on macrophage phenotype. **A)** Neutrophils were partially depleted in the critical size (4mm) muscle defect and compared to a corresponding control (fibrotic group) and the regenerative group (2mm defect). Image created with BioRender.com. **B)** Neutrophil presence in muscle at days 1 and 3. Kruskal-Wallis Test, Dunn’s post hoc. **C)** Macrophage phenotype in muscle at day 3 in the fibrotic injury group when mice were treated with anti-Ly6G antibody (Depleted group) vs. an isotype control (Isotype) or a PBS control (PBS). One-way ANOVA, Tukey’s post hoc. **D)** Hierarchical clustering of DEGs from nanoString of FACS-sorted macrophages at day 3. **E)** Representative Masson’s trichrome images of all injuries (scale bar 2 mm). **F)** Quantification of collagen staining per section. **G)** Quantification of muscle fiber diameter from laminin staining. **H)** Quantification of whole muscle cross-sectional area from Masson’s trichrome stains. One-Way ANOVA, Tukey’s post hoc. N = 8, bars show mean ± SD. * p<0.0.5; ** p < 0.01; *** p < 0.001; p < 0.0001****.

Flow cytometry analysis of macrophage phenotype at the site of injury demonstrated that neutrophil depletion increased PD-L1 expression on day 3 compared to both PBS and isotype controls (**Figure 7C, Suppl. Figure 14**). However, there were no other differences in macrophage phenotype markers between the neutrophil-depleted group and the isotype control. To further characterize macrophage phenotype following neutrophil depletion, we FACS-sorted macrophages from these injuries at day 3 and conducted gene expression analysis using nanoString (**Figure 7D, Suppl. Figure 14, Suppl. Table 5**). Analysis of individual genes showed that neutrophil depletion caused macrophages to upregulate 7 genes and to downregulate 2 genes relative to the fibrotic control (PBS) group (using a p-value cut-off of 0.01) (**Suppl. Figure 14**). Here it is important to note that results using the PBS control may differ from an isotype control.

Finally, we evaluated the effects of partial neutrophil depletion on muscle repair outcomes at day 28 using histology for total collagen staining (Masson’s trichrome) and immunohistochemistry of laminin expression as a marker of muscle fibers (**Figure 7E-H, Suppl. Figure 15**). No significant differences were observed in collagen staining, muscle fiber diameter, or muscle fiber area compared to the fibrotic control (PBS) group. Altogether, these results demonstrate that, surprisingly, partial neutrophil depletion had little to no effect on macrophage phenotype or on muscle repair outcome.

## 3. Discussion

This study demonstrated numerous changes in immune cell behavior locally within the muscle and systemically between small (regenerative) and critical size (fibrotic) VML injuries. Within 1 day after injury, macrophages exhibited very different phenotypes, especially on the gene expression level, with changes persisting for up to 4 weeks, even though overall temporal trends in typical phenotype markers were largely similar. In addition, we uncovered several differences in leukocyte numbers in the bone marrow, changes in macrophage phenotype in the spleen, and differences in inflammatory cytokine levels systemically. Lastly, we observed a large increase in neutrophil accumulation locally in fibrotic injuries compared to regenerative injuries.

Despite this finding, and a vast body of research implicating neutrophils in macrophage dysfunction, partial neutrophil depletion had little to no effect on macrophage phenotype or on muscle repair outcome. These results are critical for understanding how the dynamics of immune cell interactions differ between two injuries of different sizes in the same tissue, which lead to very different outcomes. Furthermore, they demonstrate that the entire system must be considered when designing new interventions.

The reasons for impaired tissue repair in critical sized defects are often attributed to reductions in the number of regenerative stem cells or to physical limitations of cell migration. However, our results show that dysfunctional immune cell dynamics likely also play an important role. For example, macrophage gene expression was broadly downregulated in the fibrotic injuries as early as day 1, with decreased expression of critical genes involved in regulation of repair and clearance of tissue debris. These effects would be expected to inhibit muscle regeneration even if there were sufficient levels of stem cells available for differentiation. The increased numbers of macrophages and altered phenotypes persisted to at least day 28, indicating long term consequences of the initial injury. Therefore, any strategy intended to promote muscle regeneration must take these immune system changes into account.

It is not known why the size of the injury has such drastic local and systemic effects on immune cell dynamics. It may simply be that a larger injury causes more cell injury and death (**Suppl. Figure 16**), and therefore more release of danger associated molecular patterns (DAMPs) into the circulation. Previous reports have highlighted the role of High Mobility Group Box 1 (Hmgb1), a DAMP and agonist for macrophage toll like receptor 4 (TLR4), in skewing macrophages towards an inflammatory phenotype ^39,40^. Interestingly, Hmgb1 has also been shown to decrease macrophage phagocytotic/efferocytotic capacity ^41,42^, perhaps leading to the high levels of neutrophils observed in this study. Additionally, we also observed increased expression of genes related to leukocyte-recruiting factors in the living cells remaining in the injured muscle of the fibrotic outcome groups. Together these factors could account for the increased leukocyte infiltration in the fibrotic outcome group and the altered behavior of macrophages, which are critical for regulating satellite cell proliferation and myofibroblast differentiation and myotube formation ^43^. By day 3, muscle cells had downregulated expression of key genes involved in muscle repair, including *Col1a1*, *Col5a2*, and *Dcn*. It is possible that the altered macrophages and other leukocytes drove these changes, which should be evaluated in future studies.

In many tissues, including muscle, it is believed that the transition of macrophages from pro-inflammatory (or “M1-like”) to reparative (or “M2-like”) phenotypes is critical for successful tissue regeneration, which has been reviewed by our own group ^44^ and others ^45^. However, in the present study, we found that the macrophages in both the regenerative and fibrotic outcome groups exhibited the same temporal trends of typical phenotype markers, with the only differences being in their expression levels. The fibrotic group’s macrophages expressed higher levels of CD9, CD38, and CXCR4, but these markers are not specific markers of pro-inflammatory macrophages. On the gene expression level, macrophages in the fibrotic outcome group also did not appear more pro-inflammatory than macrophages in the regenerative outcome group. They did not upregulate any of the pro-inflammatory genes in the panel, and in fact most of their differentially expressed genes were downregulated compared to the regenerative group’s macrophages. The fibrotic group’s macrophages did express lower levels of several reparative genes and proteins, including CD163, CD301b, and CD206, which are key markers of regenerative macrophages that are also involved in clearance of tissue debris. Together, these results suggest that the macrophages in the fibrotic outcome group were not overly pro-inflammatory but rather defective in their reparative/regenerative behavior. One other study that characterized macrophage phenotype in critical vs. subcritical VML also reported that macrophages from critical sized defects were no more pro-inflammatory or M1-like, based on Ly6C expression, but were actually more M2-like, based on CD206 expression, although the macrophage gating strategy differed quite a bit from the one used in the current study in that it also included the efferocytosis receptor MerTK and the Fc receptor CD64 to identify macrophages ^11^. Another major difference between that study and the present one is that that study used as 3mm defect in male mice whereas the present study used a 4mm defect in female mice, which is comparatively much more severe. Another study that included single cell RNA sequencing data from 3mm vs. 2mm defects in a combined cohort of male and female mice together with regenerative vs. fibrotic bone defects reported no obvious differences in M1-like or M2-like macrophage profiles between the two different outcomes ^16^. Together, these results show that macrophages from fibrotic VML injuries are quite different from macrophages observed in more typical models of fibrosis, such as liver and lung, in which macrophages express high levels of both M1-like and M2-like markers (reviewed in ref. ^46,47^). This difference is not entirely surprising considering fibrotic VML is characterized by too little ECM deposition, whereas liver and lung fibrosis are characterized by too much ECM deposition.

Another important finding from the present study was that there were systemic consequences of the larger, fibrotic injury that were not observed in the smaller, regenerative injury. For example, several blood cytokines were altered in the fibrotic group compared to naïve mice but not in the regenerative outcome group. The higher levels of systemic G-CSF in the fibrotic outcome group are consistent with the higher numbers of neutrophils at the site of injury, as G-CSF is crucial for neutrophil maturation. It is also noteworthy that the levels of three pro-inflammatory cytokines (IL1-beta, IL-12, and IL-15) were lower in the fibrotic outcome group at day 1. This effect may be attributed to a compensatory anti-inflammatory feedback mechanism in response to higher levels of inflammation at earlier time points, although the very early kinetics of systemic cytokines were not evaluated in this study. The fact that splenic macrophages were more inflammatory in the fibrotic outcome group compared to the regenerative group supports the hypothesis that early increases in inflammation have systemic repercussions. In theory, these effects could in turn have local influence at the site of injury. While the contribution of splenic macrophages to VML repair has not been evaluated, they have been shown to be major contributors to repair of other tissues like heart following myocardial infarction ^31^ and lung injury ^48^. Future studies should be directed towards increasing understanding of how these systemic repercussions of injury size may influence tissue repair outcome.

Defective efferocytosis (clearance of apoptotic neutrophils by macrophages) is well known to be a major source of chronic inflammation in fibrosis of multiple tissues following injury ranging from the lungs to the spinal cord ^49^. We and others have shown that efferocytosis is a potent stimulus that causes macrophages to switch from pro-inflammatory to anti-inflammatory/pro-regenerative ^18^, so impaired efferocytosis stalls this critical process of macrophage phenotype regulation. The fact that partial neutrophil depletion did not influence macrophage phenotype suggests that the elevated levels of neutrophils may result from defective macrophage behavior, not necessarily defective neutrophil behavior. In fact, impaired clearance of apoptotic neutrophils in other tissues like diabetic wounds has also been linked to defective macrophage activity ^50^. In this study, we did find that macrophages from the fibrotic group expressed higher levels of some pro-inflammatory phenotype markers, which would be expected to reduce their efferocytotic behavior ^21,51^, as well as decreased expression of phagocytic and efferocytotic genes at the early stages of the response to injury in this study. Neutrophils release pro-inflammatory factors, so their persistence in the injury site can impair satellite cell differentiation and fusion into myofibers ^14^. Additionally, apoptotic neutrophils that are not adequately cleared may further progress to secondary necrosis/NETosis ^52^. These modes of cell death are more inflammatory compared to apoptosis and have been shown to further impede phagocytosis by macrophages ^53^.

The present study also showed that clearance of neutrophils alone was insufficient to improve muscle repair outcomes. In fact, there have been three other published studies in which neutrophils were depleted in the context of muscle injury, and they yielded conflicting results. Sadtler et al. reported no effect of neutrophil depletion on biomaterial-mediated fibrosis ^54^. Two other studies depleted neutrophils in muscle repair using models in which injury was induced by injection of snake venom ^55^ or by exhaustive exercise ^56^, and these studies reported improved muscle repair outcomes. Therefore, more studies are required to address the lack of consensus of the effects of neutrophil depletion on muscle repair outcomes.

Although this study uncovered several previously unknown phenomena in VML repair and pathology, there were several limitations. The animal model utilized a relatively sterile injury, which is not normally the case for most VML patients. It is likely that during a non-sterile injury such as a vehicle accident or combat wounds, the elevated presence of neutrophils is crucial to clear the tissue from bacteria and other microbes. Additionally, VML in humans is often accompanied by substantial injury to the nearby bone, which can influence VML healing ^57^. While we reported some differences in neutrophil phenotype between regenerative and fibrotic outcomes, these results need to be thoroughly validated because the tissue digestion process can influence neutrophil phenotype. Furthermore, neutrophil depletion was accomplished via systemic injection of anti-Ly6G antibodies, which may depend on clearance by splenic and liver macrophages ^58,59^. Therefore, in principle this method could indirectly modulate muscle macrophage phenotype. Finally, the study was performed in female mice. Further studies must be conducted to evaluate the influence of sex on the findings described here.

In this study we have contributed to the growing body of literature concerning immune cell dysfunction following VML by identifying local and systemic immune cell differences in regenerative vs. fibrotic outcomes. Additional studies are required to fully evaluate the mechanisms involved in these systemic changes, as well as any possible feedback on immune behavior locally in the muscle tissue.

## Conclusions

This study increases understanding of macrophage dysfunction following VML by establishing a complete timeline in the context of the complex ecosystem in which they operate. Coinciding with alterations in macrophage phenotype were changes in numerous other leukocytes in the muscle and in the bone marrow, changes in splenic macrophage phenotype, and changes in systemic levels of pro-inflammatory cytokines. Despite excessive levels of apoptotic neutrophils in the muscle, their depletion to levels observed in the regenerative outcome group did not influence macrophage phenotype or muscle repair outcome. Together, these results suggest that the entire system of immune cell interactions will need to be considered to improve muscle repair outcomes in VML.

## METHODS

### Animal Surgeries

All animal experiments were conducted with approval by the Drexel University Institutional Animal Care and Use Committee. For each experiment, six to eight female C57BL/6 mice per group per time point were anesthetized using isoflurane (Vedco) and weighed. After buprenorphine sustain release pain relief administration at 0.05 mg/kg, fur was removed using depilatory cream (Nair) and the skin was washed. Next, a triple alternating iodine and ethanol treatment was used to sterilize the skin and a 1-1.5 cm incision was made in the left hindlimb. Using a 2 or 4 mm in diameter biopsy punches full-thickness injuries were created in the mid portion of the quadriceps, with the leg comfortably extended. Hemostasis was accomplished using sterile cotton tipped swabs until bleeding ceased. Animals were sutured using a number 6 suture and observed until fully recovered.

### Bone marrow derived macrophage cell culture

Femur and tibia of female C57BL/6 mice, 8-12 weeks of age, were harvested for bone marrow extraction. The ends of the bones were removed using a scalpel, and marrow was flushed using a 27 gauge containing sterile RPMI 1640 (GIBCO CAT:11875-093). Cells were then cultured in non-tissue culture treated flasks (Thermo Scientific, CAT: 12-566-85) with media containing 89% RPMI 1640, 10% Fetal Bovine Serum (FBS) (VWR, CAT: 45000-734) and 1% Penicillin-Streptomycin (Themo Fisher, CAT: 15070063) and 25 ng/ml of M-CSF (Peprotech, CAT: 315-02). Media was changed after 3 days, and 2 days later cells were detached using 3 ml of TrypLE (Thermo Fisher, CAT: 12604021). Cells were moved to a non-tissue culture treated 6-well plate (VWR, CAT: 351146) and allowed to rest overnight. Next, cells were polarized using either 40 ng/ml of IL-4 (Peprotech, CAT: 214-14), 50 ng/ml of LPS (Fisher Scientific, CAT: NC9039780) or 60 ng/ml of IL-17A (Peprotech, CAT: 210-17). Cells were collected and analyzed via flow cytometry 3 days later.

### Neutrophil depletion

Partial neutrophil depletion was accomplished via tail vein injection of anti-Ly6G antibodies (BioXCell CAT: BE0075-1) compared to a corresponding isotype control or PBS control. The antibody was diluted to 0.5 mg/ml in 100 uL of sterile PBS, and administered immediately prior to surgery, as well as 1 and 2 days after surgery. Partial neutrophil depletion was confirmed in muscle via flow cytometry, as described below.

### Tissue Preparation

For flow cytometry analysis, tissues were excised and digested into homogenous single cell suspensions. Whole muscle was digested using 3 mg/ml of Collagenase 4 (Worthington CAT:LS004188) diluted in RPMI (GIBCO CAT:11875-093) to a total volume of 3 ml for 30 minutes at 37C. Spleen was gently dissociated by pressing it against a 70-micron filter using a syringe plunger, followed by the addition of 7 ml of red blood cell (RBC) lysis (eBioscience CAT: 00-4300-54)) for 5 minutes at room temperature. Bone marrow was collected by removing the ends of both tibia and femur using a scalpel, then flushed through with ice cold RPMI using a 21 G syringe. The marrow was then lysed using the same protocol as the spleen. Blood was collected via cardiac puncture using a 25G needle and placed in an EDTA coated tube (BD CAT: 367856). Blood was then centrifuged at 2,000 G for 5 minutes to separate the serum from the cells. The serum was frozen at –80C for Luminex, while the cell pellet underwent two rounds of RBC lysis as previously described.

### Flow Cytometry

All samples were plated at approximately 1 million cells per well, where appropriate Fluorescence Minus One (FMOs) and Unstained controls were created by mixing cells from all samples, where each organ had its own FMO. Staining dilutions, catalog numbers and vendors can be found in **Table 1**. Cells were first stained with 200 uL Live/Dead for 30 minutes at 4C. Next, cells were stained with 50 uL FcR block for 10 minutes at 4C. Cells then received 50 uL of extracellular marker staining for 15 minutes at 4C. Next, samples were washed and fixed using BD Fixation and Permeation buffer for 20 minutes at 4C. Cells were washed twice with the appropriate washing buffer and stained for intracellular markers for 30 minutes at room temperature. Finally, cells were washed and resuspended in FACS buffer at 4C until analysis.

**Table 1.** List of antibodies.

| Marker | Color | Catalog | Dilution | Vendor |
| --- | --- | --- | --- | --- |
| CD45 | FITC | 147710 | 1:100 | Biolegend |
| Ly6G | PeCy7 | 127618 | 1:75 | Biolegend |
| Ly6G | APC-Cy7 | 127624 | 1:75 | Biolegend |
| Ly6C | APC | 128015 | 1:75 | Biolegend |
| CD3 | Brilliant Violet 605 | 100237 | 1:50 | Biolegend |
| CD11c | PeDazzle 594 | 117348 | 1:75 | Biolegend |
| NK1.1 | Alexa Fluor 700 | 156512 | 1:75 | Biolegend |
| B220 | Brilliant Violet 421 | 103240 | 1:60 | Biolegend |
| F4/80 | APC-Cy7 | 123118 | 1:50 | Biolegend |
| F4/80 | Brilliant Violet 711 | 123147 | 1:50 | Biolegend |
| CD11b | Pe | 101208 | 1:75 | Biolegend |
| CD9 | PerCp-Cy5.5 | 124818 | 1:50 | Biolegend |
| CD38 | Alexa Fluor 700 | 102742 | 1:100 | Biolegend |
| CD86 | Brilliant Violet 785 | 105043 | 1:75 | Biolegend |
| PD-L1 | PeDazzle 594 | 124324 | 1:50 | Biolegend |
| CXCR4 | Brilliant Violet<br>605 | 146519 | 1:50 | Biolegend |
| Arg1 | APC | 17-3697-82 | 1:50 | Invitrogen |
| CD163 | Pe | 155308 | 1:75 | Biolegend |
| CD301b | PeCy7 | 146808 | 1:50 | Biolegend |
| CD206 | Brilliant Violet<br>421 | 141717 | 1:50 | Biolegend |
| CD16/32 (FcR<br>Block) | N/A | 156604 | 1:200 | Biolegend |
| Live/Dead Aqua | Brilliant Violet<br>510 | L34966 | 1:1,000 | Invitrogen |
| ApoTracker | Green (FITC) | 427403 | 1:1,600 | Biolegend |

**Table 2.** Gating Strategy.

| **Gating Strategy** | **Immune Cell** |
| --- | --- |
| Live/CD45+/B220+ | B Cells |
| Live/CD45+/CD3+ | T Cells |
| Live/CD45+/NK1.1+ | Natural Killer Cells |
| Live/CD45+/B220-/CD3-/NK1.1-/CD11b+/Ly6G-/CD11c+ | Dendritic Cells |
| Live/CD45+/B220-/CD3-/NK1.1-/CD11b+/Ly6G+ | Neutrophils |
| Live/CD45+/B220-/CD3-/NK1.1-/CD11b+/Ly6G-/CD11c-<br>/Ly6C High | Classical Monocytes |
| Live/CD45+/B220-/CD3-/NK1.1-/CD11b+/Ly6G-/CD11c-<br>/Ly6C Low | Non-classical Monocytes |
| Live/CD45+/B220-/CD3-/NK1.1-/Ly6G-/F480+ | Macrophages |

Samples destined for Fluorescence Activated Cell Sorting (FACS) were prepared similarly to samples prepared for flow cytometry analysis, except samples were split in half. One half was used for sorting and follow up RNA extraction, while the other half was used to create the appropriate FMO and unstained control. The volume of blocking and staining cocktail was adjusted accordingly, maintaining the same dilution. Note that F4/80 was not used in the gating strategy as F4/80 expression has been previously described in immature neutrophils ^60^. Moreover, we confirmed the presence of a small subpopulation of F4/80+ neutrophils that did not differ between both groups (**Suppl. Figure 10**). All samples were analyzed using a BD Fortessa. Sample analysis was conducted using FlowJo. Clustering analysis was performed by identifying the number of clusters present using XShift, followed by clustering using FlowSOM using the number of clusters from the previous step.

### nanoString multiplex gene expression analysis

RNA was extracted using RNAqueous microkit (Invitrogen CAT: AM1982) for FACS-sorted macrophages and neutrophils, while whole tissue RNA was extracted using RNAqueous isolation kit (Invitrogen CAT: AM1912), according to manufacturer’s recommendation. RNA was evaluated for quantity and purity using Biotek Take3^TM^ plate and Biotek synergy H1 microplate reader. 5 uL of 20 ng/ul per sample was added to the reaction tube. Next 8 uL of master mix was added to the sides of each tube, followed by 2 uL of Capture Probeset. Samples were quickly centrifuged to allow reagents to mix and placed on thermocycler at 65C for 20 hours. Gene counts were measured using nCounter MAX/FLEX system, and samples were scanned using the high field of view (FOV, 280) settings.

### Histology and immunohistochemistry (IHC)

Histopathological and IHC assays were performed by Translational Pathology Shared Resources at Sidney Kimmel Cancer center, Thomas Jefferson University. Mouse quadriceps muscle tissue samples were harvested and fixed in 10% Neutral Formalin Buffer (NFB) fixative at room temperature for 48 hours and then embedded in paraffin. These formalin fixed paraffin embedded (FFPE) blocks were sectioned at 4 µm. Four levels with 1500 µm apart were examined beginning from distal part of the muscle. H&E and Masson’s Trichrome staining were performed by using standard methods. IHC staining was performed using an intelliPATH FLX® Automated Slide Stainer (Biocare medical, LLC). Antibodies and epitope retrieval methods are: PECAM-1/CD31 (Santa Cruz, Cat#: sc-1506-R, 1/200 dilution), Heat Induced Epitope Retrieval (HIER) using Uni-TRIEVE (INNOVEX, Cat#: NB325-500) at 70°C for 1 hour; Laminin (Abcam, Cat#: ab11575, 1/400 dilution), epitope retrieval was performed using proteinase K(Invitrogen, P/N: 100005393, 20µg/ml) at room temp for 5 minutes. Briefly, the slides were deparaffinized, rehydrated and then exposed to epitope retrieval. After endogenous peroxidase, normal serum and endogenous biotin blocking, primary antibodies were applied and incubated at room temperature. Biotinylated anti-rabbit (Vector Laboratories, cat#: BA-1000) secondary antibody and ABC-HRP complexes (Vector Laboratories, Cat#: PK-6100) were applied following the primary antibodies with 30 minutes incubation at room temperature. Three washes with 1x tris-buffered saline with Tween (TBTS) were performed between each step above. The signals were visualized using ImmPACT™ VECTOR Red (Vector Laboratories, Cat#: SK5105). The slides were counter stained with Hematoxylin, dehydrated, cleared and coverslipped with Permount mounting medium. All the stained slides were scanned using Aperio ScanScope CS2 at 20X magnification.

### Luminex

Samples were analyzed using manufacturer’s recommendation. Blood was collected via cardiac puncture using a 25G needle and placed in an EDTA coated tube (BD CAT: 367856). Blood was then centrifuged at 2,000 G for 5 minutes to separate the serum from the cells. The serum was then frozen at –80C for Luminex.

### Histological analysis and sGAG quantification

Collagen accumulation was measured as a function of blue pixels (collagen) in Masson’s Trichrome staining over all pixels using a custom MATLAB code (**Suppl. Methods**). Muscle fiber diameter was measured manually using Laminin stained sections. Using ImageScope software, 4 different regions of interest surrounding the center of the injury (below, right and left), including the center, were selected and 50 measurements were taken at random per region, totaling 200 measurements per sample. Only one section per sample was used in this quantification. These values were averaged resulting in a single average muscle fiber diameter per sample. Similarly, muscle cross sectional area was measured using ImageScope software by contouring the outer edge of the muscle section. sGAG content was measured by first digesting tissue with 2% papain (Millipore Sigma, CAT:76216) (1 mL/construct, 0.56 U/mL in a buffer containing 0.1 M sodium acetate (Millipore Sigma, CAT: 71188), 10 M cysteine hydrochloric acid (Millipore Sigma, CAT: 30120), 0.05 M EDTA (MP Biomedicals, CAT: 6381-926), pH 6.0) at 60 °C for 16 h. Sulfated glycosaminoglycans (sGAG) content was quantified with DMMB assay following established procedures ^61^. These data were normalized by the wet weight of the samples for native tissues (N ≥ 4 animals).

### Statistical Analysis

Statistical analysis of flow cytometry data and quantification of histological images was performed using GraphPad Prism v10. All data were reported as mean values ± standard deviation. Outliers were detected using ROUT method, and removed if deemed an outlier. The normality of data distributions was verified via Kolmogorov-Smirnov test. Depending on normality, data were analyzed with one-way or two-way ANOVA or respective non-parametric test or multiple t-tests, as specified in each figure legend. All measurements reported were taken from distinct samples, and not repeated measurement. Where applicable, two-sided tests were applied. The z-score for each marker was calculated based on the expression of that marker across all clusters using weighted means and standard deviation, to account for the discrepancy of cluster sizes.

NanoString data were normalized according to manufacturer recommendations. Briefly, negative control was first subtracted from all gene counts, followed by a positive control normalization. Then, gene counts were normalized by total counts per sample.

Negative values were removed, and all data was transformed on log base 10. For statistical analysis, data were log-transformed and multiple t-tests were conducted with a p<0.01 threshold. This method was selected due to the interdependency of each gene, which does not allow for conventional multiple comparisons’ corrections, so a more conservative p-value was selected to limit the number of type 1 errors. Clustering analysis of nanoString data was performed using RStudio (R Version 4.3.1). The normalized and log transformed gene counts were converted into Z Scores, and clustering was done by minimizing cluster variance.

## Data Availability

All data is available upon request from the corresponding author.

## Code Availability

The custom MATLAB code used to quantify collagen staining from Masson’s trichrome images is included in the Supplementary Material.

## Acknowledgments

This study was supported by NHLBI R01 HL130037 and NIAMS R21 AR083080 to KLS. We would like to acknowledge the following Shared Resources at Sidney Kimmel Cancer center, Thomas Jefferson University that were used to conduct this research: Translational Pathology Shared Resources (supported by the NCI grant 5P30CA056036-23). The authors thank Drs. Yangzhu Du, Honghong Sun, and Eline Luning Prak of the Human Immunology Core (HIC) at the Perelman School of Medicine at the University of Pennsylvania for assistance with Luminex assay. The HIC is supported in part by NIH P30 AI045008 and P30 CA016520. HIC RRID: SCR_022380. Additionally, we would like to acknowledge the SKCC Flow Cytometry and Cell Sorting Facility, supported by NCI Core grant (P30 CA056036).

## Author Contributions

RW and KLS designed the study and the experiments. LH, BBM, and RJP contributed intellectually to the design of experiments and interpretation of results. SS, GER, PEC, EMO, TT, and TL contributed to experimental execution and data acquisition. All authors have read and approved the manuscript.

## Competing Interests

The authors declare no competing interests.

## Notes

### Competing Interest Statement

The authors have declared no competing interest.

