## Supplemental Material for "Effects of injury size on local and systemic immune cell dynamics in volumetric muscle loss"

Whitaker et al.

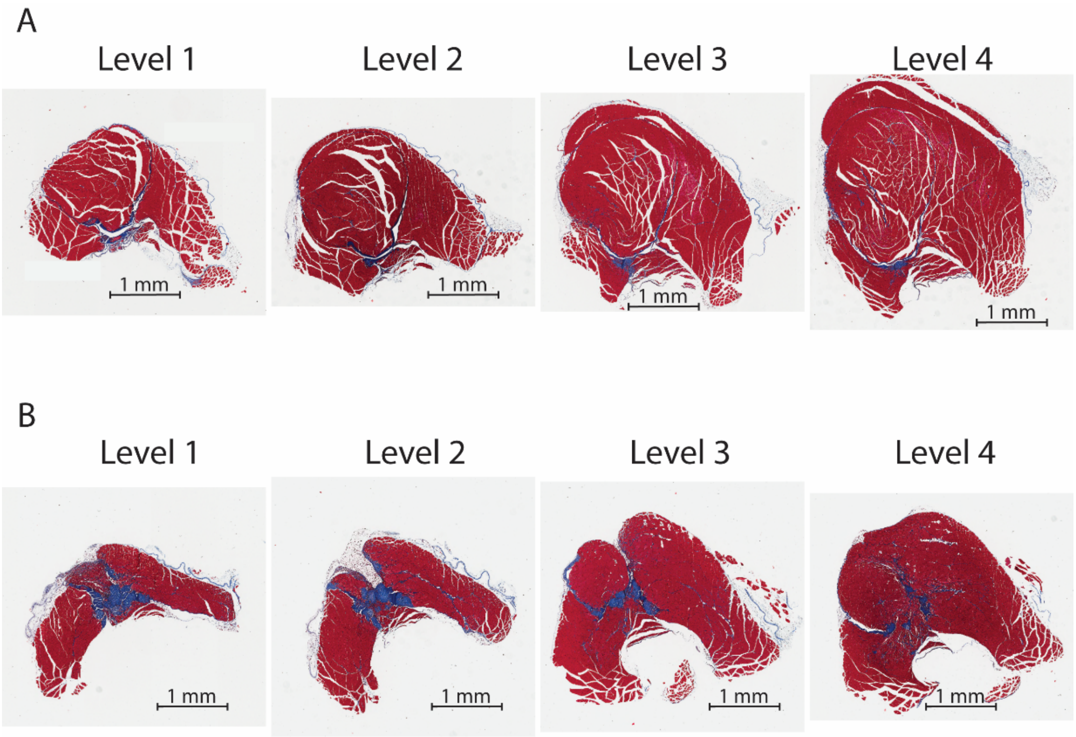

**Suppl. Figure 1.** Muscle sections stretching 6 mm along the center of muscle injury. Level 1 initiates 3 mm from the distal portion of the muscle, and each level is 1.5 mm apart from the previous one. **A)** Regenerative. **B)** Fibrotic.

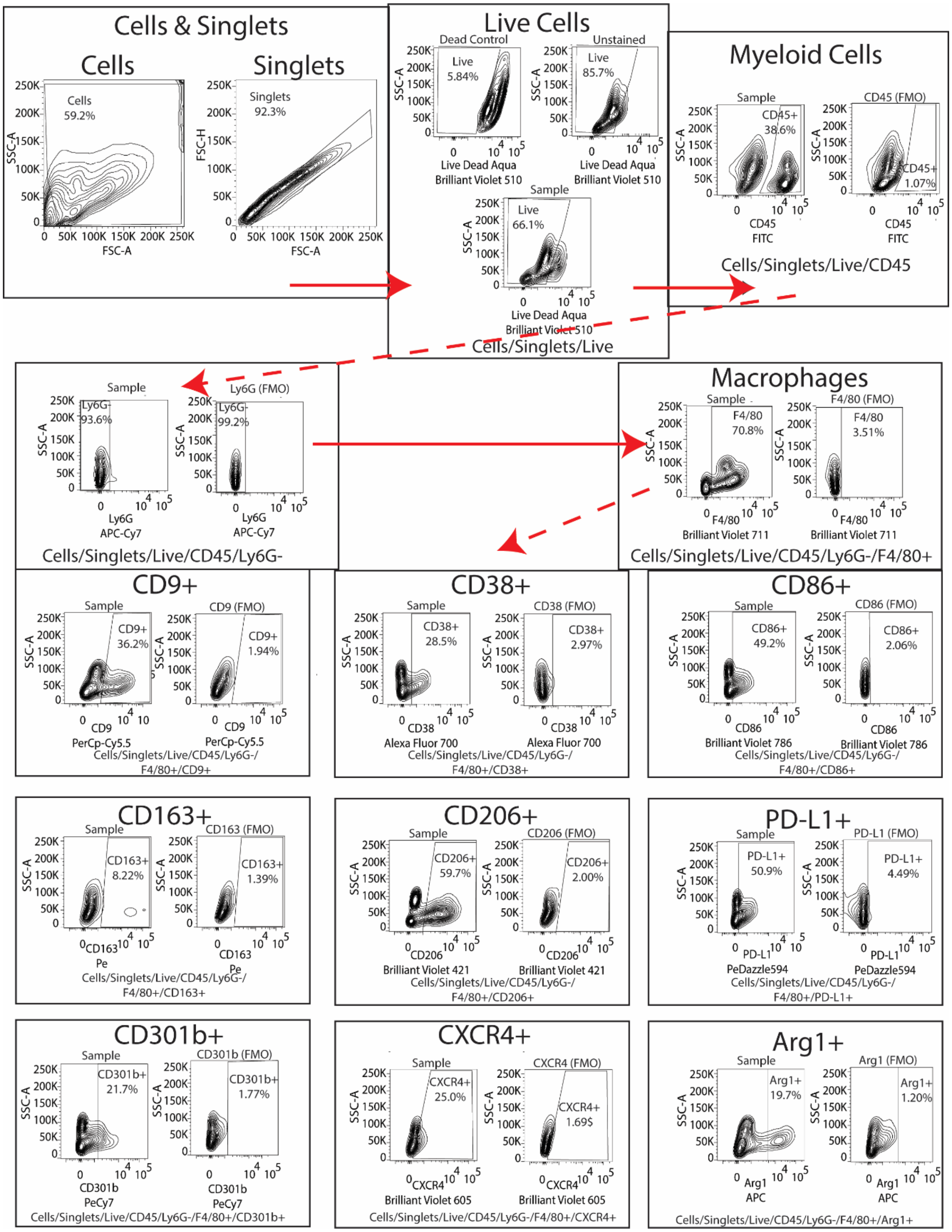

**Suppl. Figure 2.** Macrophage phenotype gating strategy via flow cytometry.

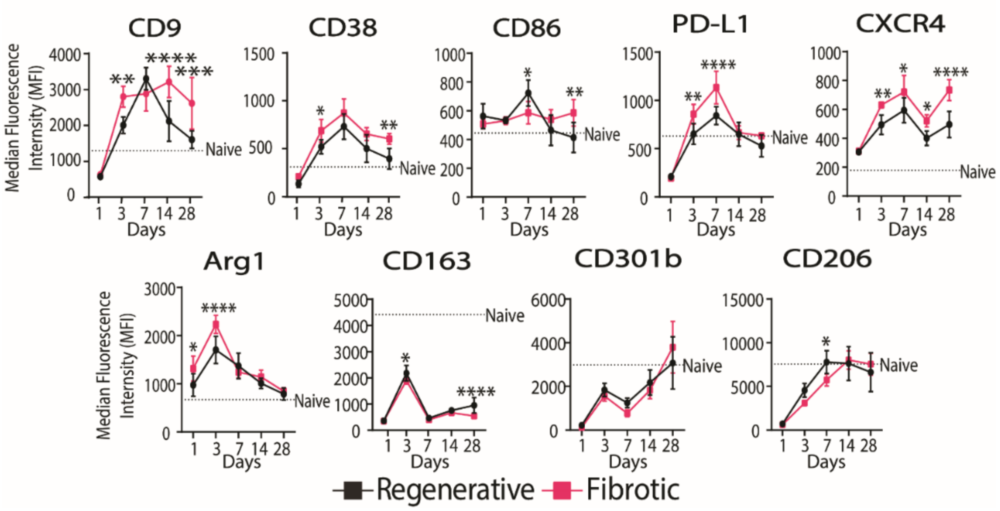

**Suppl. Figure 3.** Muscle macrophage phenotype measured by flow cytometry. Two-Way ANOVA, Sidak's Post Hoc. N = 6, bars show mean ± SD. p<0.0.5 * ; p < 0.01 ** ; p < 0.001 *** ; p < 0.0001 ****

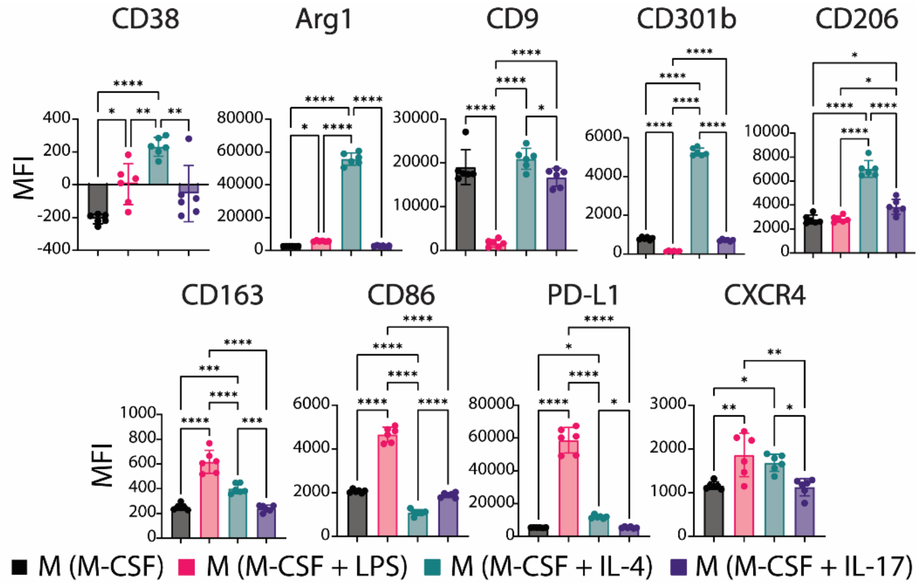

**Suppl. Figure 4.** Flow cytometry analysis of pre-polarized in vitro macrophage phenotype.

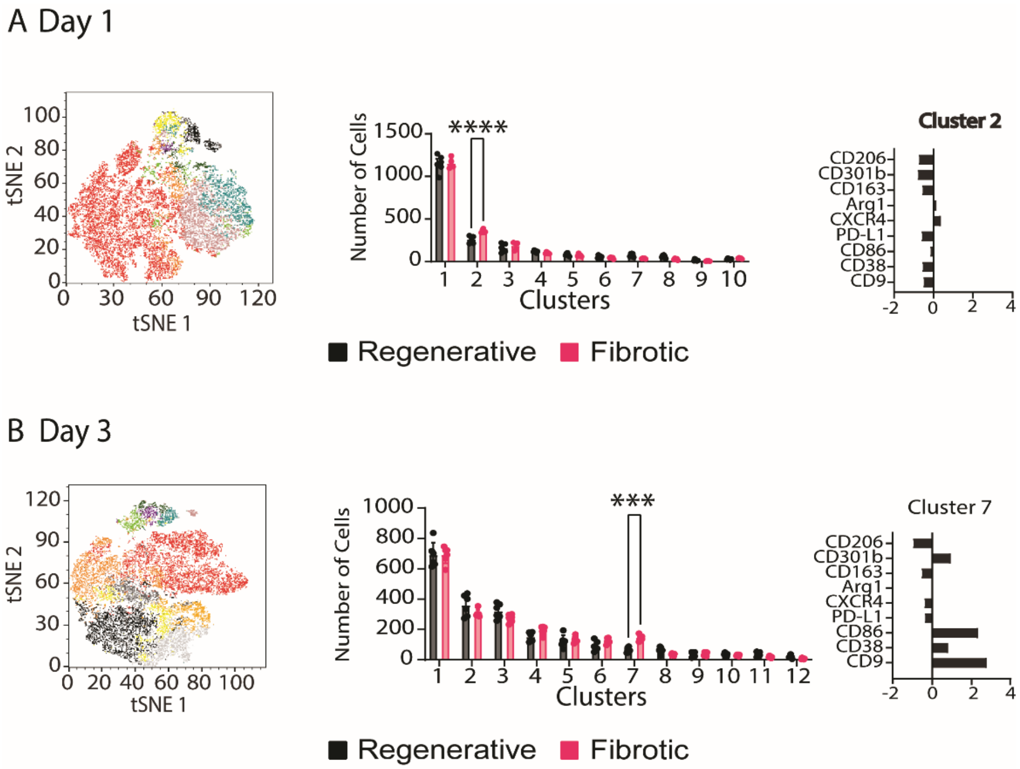

**Suppl. Figure 5. A)** Hierarchical clustering of MFI values at Day 1. **B)** Hierarchical clustering of MFI values at Day 3. Two-Way ANOVA, Sidak's Post Hoc. N = 6, bars show mean ± SD. p<0.0.5 * ; p < 0.01 ** ; p < 0.001 *** ; p < 0.0001 ****

**Suppl. Table 1.** List of genes and category in NanoString panel

| **Gene Name** | **Category** | **Gene Name** | **Category** | **Gene Name** | **Category** | **Gene Name** | **Category** |
|---|---|---|---|---|---|---|---|
| *Acta2* | ECM | *Cltc* | Phagocytosis | *Il18* | Inflammatory | *Psme1* | Immune Response |
| Agtr1a | Inflammatory | *Col1a1* | ECM | *IL1a* | Inflammatory | *Psme2* | Immune Response |
| *Angpt1* | Angiogenesis | *Col1a2* | ECM | *Il1b* | Inflammatory | *Ager* | Inflammatory |
| *Angpt2* | Angiogenesis | *Col3a1* | ECM | *Il2* | Immune Response | *Rela* | Immune Response |
| *Angpt4* | Angiogenesis | *Col5a1* | ECM | *Il23r* | Immune Response | *Relb* | Immune Response |
| *Anxa1* | Phagocytosis | *Col5a2* | ECM | *Il36g* | Immune Response | *Retnla* | Reparative |
| *Ap2a2* | Phagocytosis | *Col6a1* | ECM | *Il4* | Macrophage activation | *Rnase2a* | Immune Response |
| *Apoe* | Phagocytosis | *Cox5a* | Immune Response | *Il4ra* | Reparative | *S100A4* | Immune Response |
| *Areg* | Inflammatory | *Rel* | Immune Response | *Il6* | Inflammatory | *S1pr1* | Immune Response |

| | | | | | | | |
|---|---|---|---|---|---|---|---|
| *Arg1* | Reparative | *Csf1* | Immune Response | *Irf4* | Reparative | *S1pr2* | Immune Response |
| *Asprv1* | Immune Response | *Csf1r* | Immune Response | *Irf5* | Inflammatory | *S1pr3* | Immune Response |
| *Axl* | Phagocytos is | *Cst7* | Immune Response | *Irf7* | Immune Response | *S1pr4* | Immune Response |
| *Bgn* | ECM | *Cstsb* | Immune Response | *Isg15* | Phagocytosis | *S1pr5* | Immune Response |
| *Bmp2* | Trafficking | *Ctsd* | Immune Response | *Itgax* | Inflammatory | *Selenop* | Reparative |
| *C3ar1* | Trafficking | *Ctsl* | Immune Response | *Itgb5* | Trafficking | *Socs3* | Phagocytosis |
| *C5ar1* | Trafficking | *Ctsz* | Immune Response | *Jag1* | Angiogenesis | *Spp1* | ECM |
| *Cbr2* | Reparative | *Cx3xl1* | Trafficking | *Jak1* | Immune Response | *Stat1* | Inflammatory |
| *Ccl1* | Reparative | *Cx3cr1* | Trafficking | *Jak2* | Immune Response | *Stat2* | Immune Response |
| *Ccl12* | Inflammato ry | *Cxcl1* | Trafficking | *Jak3* | Immune Response | *Stat3* | Reparative |
| *Ccl17* | Reparative | *Cxcl10* | Trafficking | *Klf4* | Reparative | *Stat4* | Immune Response |
| *Ccl2* | Trafficking | *Cxcl11* | Trafficking | *Lag3* | Immune Response | *Stat5a* | Immune Response |
| *Ccl22* | Reparative | *Cxcl12* | Trafficking | *Lpl* | Phagocytosis | *Stat5b* | Immune Response |
| *Ccl24* | Reparative | *Cxcl2* | Trafficking | *Sell* | Trafficking | *Stat6* | Reparative |
| *Ccl4* | Trafficking | *Cxcl3* | Trafficking | *Nr1h3* | Immune Response | *Stx5a* | Houskeeping gene |
| *Ccl5* | Trafficking | *Cxcr2* | Trafficking | *Nr1h2* | Immune Response | *Tbp* | Houskeeping gene |
| *Ccl6* | Trafficking | *Cxcr3* | Trafficking | *Itgb7* | Trafficking | *Tek* | Angiogenesis |
| *Ccl7* | Trafficking | *Cxcr4* | Trafficking | *Ly6c1* | Immune Response | *Tgfb1* | ECM |
| *Ccl8* | Trafficking | *Dcn* | ECM | *Itgam* | Trafficking | *Tgfb2* | Immune Response |
| *Ccn2* | ECM | *Egf* | ECM | *Mapk1* | Immune Response | *Tgfb3* | Immune Response |
| *Ccr2* | Inflammato ry | *Egr2* | Reparative | *Marco* | Reparative | *Tgfbr2* | Immune Response |
| *Ccr4* | Trafficking | *Eno1* | Immune Response | *Ly96* | Inflammatory | *Tgfbr3* | Immune Response |
| *Ccr5* | Trafficking | *Sele* | Immune Response | *Mertk* | Phagocytosis | *Tie1* | Angiogenesis |
| *Ccr6* | Trafficking | *Adgre1* | Trafficking | *Mki67* | Immune Response | *Timp1* | ECM |
| *Ccr7* | Trafficking | Fap | ECM | *Mmp10* | Inflammatory | *Timp2* | ECM |
| *Ccr8* | Trafficking | *Fcgrt* | Phagocytos is | *Mmp13* | ECM | *Timp3* | ECM |
| *Itgb4* | Immune Response | *Fgf1* | Immune Response | *Mmp2* | Inflammatory | *Tlr1* | Immune Response |
| *Itgal* | Trafficking | *Fgf2* | Immune Response | *Mmp7* | ECM | *Tlr11* | Immune Response |

| | | | | | | | |
|---|---|---|---|---|---|---|---|
| *Cd14* | Inflammatory | *Fgf7* | Immune Response | *Mmp8* | ECM | *Tlr2* | Immune Response |
| *Ctla4* | Immune Response | *Flt1* | Angiogenesis | *Mmp9* | Reparative | *Tlr3* | Immune Response |
| *Cd163* | Reparative | *Fn1* | ECM | *Mrc1* | Reparative | *Tlr4* | Immune Response |
| *Itgb2* | Trafficking | *Foxp3* | Immune Response | *MyD88* | Inflammatory | *Tlr5* | Immune Response |
| *Il2ra* | Immune Response | *Gusb* | Houskeeping gene | *Ndufa1* | Immune Response | *Tlr6* | Immune Response |
| *Itgb1* | Trafficking | *H2-Aa* | Immune Response | *Ndufc2* | Immune Response | *Tlr7* | Immune Response |
| *Mgl2* | Reparative | *H2-Ab1* | Immune Response | *Nfkb1* | Inflammatory | *Tlr8* | Immune Response |
| *Cd36* | Inflammatory | *H2-DMa* | Immune Response | *Nfkb2* | Inflammatory | *Tlr9* | Immune Response |
| *Cd38* | Inflammatory | *H2-DMb1* | Immune Response | *Nfkbia* | Inflammatory | *Tme* | Immune Response |
| *Cd3e* | Immune Response | *H2-Eb1* | Immune Response | *Nfkbiz* | Immune Response | *Tnf* | Inflammatory |
| *Cd4* | Immune Response | *Hgf* | Immune Response | *Ngf* | Immune Response | *Tmem119* | Immune Response |
| *Cd52* | Immune Response | *Hif1a* | Immune Response | *Nlrp3* | Inflammatory | *Trem2* | Immune Response |
| *Itgb3* | Trafficking | *Hnrnpab* | Houskeeping gene | *Nos2* | Inflammatory | *Ticam2* | Immune Response |
| *Cd63* | Phagocytosis | *Hspg2* | ECM | *P2ry12* | Immune Response | *Txnip* | Phagocytosis |
| *Fcgr1* | Inflammatory | *Iba1* | Phagocytosis | *P2ry13* | Immune Response | *Tyrobp* | Immune Response |
| *Cd68* | Inflammatory | *Ifna* | Macrophage activation | *Pcna* | Immune Response | *Ubb* | Immune Response |
| *Cd69* | Immune Response | *Ifna1* | Macrophage activation | *Pdgfb* | Angiogenesis | *Ubc* | Immune Response |
| *Tfrc* | Immune Response | *Ifnb1* | Macrophage activation | *Pdgfra* | Immune Response | *Uqcrq* | Immune Response |
| *Cd74* | Inflammatory | *Ifng* | Macrophage activation | *Pdgrfb* | Immune Response | *Vcan* | ECM |
| *Cd80* | Inflammatory | *Igf1* | Reparative | *Cd274* | Inflammatory | *Vegfa* | Angiogenesis |
| *Cd83* | Inflammatory | *Igf2* | Reparative | *Pdcd1lg2* | Reparative | *Vim* | ECM |
| *Cd86* | Inflammatory | *Il10* | Reparative | *Pecam1* | Angiogenesis | *Itga4* | ECM |
| *Cd8a* | Immune Response | *Il11* | Immune Response | *Pigf* | Angiogenesis | *Wfdc17* | Reparative |
| *Cd9* | Inflammatory | *Il12a* | Inflammatory | *Plod2* | ECM | *Wfdc21* | Inflammatory |
| *Chil3* | Reparative | *Il12b* | Inflammatory | *Pparg* | Reparative | *Wnt-0b* | Reparative |
| *Clec10a* | Reparative | *Il13* | Immune Response | *Selplg* | Immune Response | *Wnt3a* | Immune Response |
| *Clecl7a* | Reparative | *Il17a* | Immune Response | *Psma3* | Immune Response | | |

| | | | | | |
|---|---|---|---|---|---|
| *Clta* | Phagocytosis | *Il17ra* | Immune Response | *Psma4* | Immune Response |

**Suppl. Table 2.** List of differentially expressed genes (DEGs) in FACS-sorted macrophages, with p-value and fold change (Fibrotic vs. Regenerative).

| Day 1 | | | Day 3 | | |
|---|---|---|---|---|---|
| **Gene Name** | **P Value** | **Fold Change** | **Gene Name** | **P Value** | **Fold Change** |
| *Itgb1* | 0.000041 | -0.08396 | *Cd83* | 0.00006 | -0.1507 |
| *Ly96* | 0.000052 | -0.07572 | *Mgl2* | 0.000303 | -0.3643 |
| *Adgre1* | 0.000219 | -0.1175 | *Apoe* | 0.000884 | -0.09443 |
| *Clta* | 0.000244 | -0.07949 | *Adgre1* | 0.000933 | -0.0847 |
| *Cxcl2* | 0.000351 | 0.08269 | *Jak1* | 0.001484 | -0.09948 |
| *Cox5a* | 0.000522 | -0.06448 | *Ctsl* | 0.001739 | 0.07381 |
| *Apoe* | 0.000553 | -0.1549 | *Cd52* | 0.001747 | 0.1311 |
| *Selenop* | 0.000702 | -0.2231 | *Timp2* | 0.002081 | -0.07908 |
| *Gusb* | 0.000945 | -0.08402 | *Itgal* | 0.002221 | 0.1762 |
| *Cbr2* | 0.000957 | -0.2329 | *Retnla* | 0.002304 | -0.6417 |
| *Cd83* | 0.000986 | -0.07861 | *Selplg* | 0.002362 | -0.07666 |
| *Csf1* | 0.001117 | 0.2985 | *Egr2* | 0.002425 | 0.1751 |
| *Tgfbr2* | 0.001379 | -0.1535 | *Txnip* | 0.002596 | -0.09388 |
| *Cltc* | 0.00138 | -0.07193 | *Mmp8* | 0.002612 | 0.4208 |
| *Dcn* | 0.001411 | -0.5593 | *Ly96* | 0.003842 | -0.1094 |
| *Mapk1* | 0.001433 | -0.06362 | *C3ar* | 0.004385 | -0.08364 |
| *Cxcl3* | 0.001752 | 0.2626 | *Ccr2* | 0.004498 | -0.08321 |
| *Col3a1* | 0.001779 | -0.5239 | *Trem1* | 0.005157 | 0.3528 |
| *Uqcrq* | 0.002561 | -0.07327 | *Stat3* | 0.005334 | -0.08612 |
| *Jak1* | 0.002599 | -0.05427 | *Csf1r* | 0.006187 | -0.07911 |
| *Timp2* | 0.002892 | -0.2304 | *Cbr2* | 0.008032 | -0.1791 |
| *Ap2a2* | 0.002985 | -0.07011 | *Selenop* | 0.008274 | -0.1409 |
| *Ubb* | 0.003263 | -0.02238 | *Itgb1* | 0.008507 | -0.06463 |
| *Tfrc* | 0.003275 | -0.1498 | | | |
| *Ctsz* | 0.003574 | -0.03727 | | | |
| *Ctsb* | 0.004836 | -0.08792 | | | |
| *Clec10a* | 0.005236 | -0.2669 | | | |
| *Stat5b* | 0.005335 | -0.07517 | | | |
| *Cx3cr1* | 0.005498 | -0.2638 | | | |
| *Tlr4* | 0.006005 | -0.05434 | | | |
| *Fcgrt* | 0.00675 | -0.1733 | | | |
| *Itga4* | 0.007284 | -0.1248 | | | |
| *Selplg* | 0.008753 | -0.09944 | | | |
| *Cd68* | 0.009191 | -0.04641 | | | |

| | | |
|---|---|---|
| *Hnrnpab* | 0.009769 | -0.05098 |

**Suppl. Table 3.** List of differentially expressed genes (DEGs) in whole tissue, with p-value and fold change (Fibrotic vs. Regenerative).

| 6 Hours | | | Day 1 | | | Day 3 | | |
|---|---|---|---|---|---|---|---|---|
| **Gene Name** | **P Value** | **Fold Change** | **Gene Name** | **P Value** | **Fold Change** | **Gene Name** | **P Value** | **Fold Change** |
| *Egf* | 0.000003 | -0.9124 | *Cxcl2* | 0.000004 | 0.2123 | *Col5a2* | <0.000001 | -0.1122 |
| *Il17ra* | 0.000017 | 0.1534 | *C5ar* | 0.000115 | 0.09327 | *Ctsd* | 0.000003 | 0.1775 |
| *Tlr6* | 0.00002 | 0.2993 | *Igf1* | 0.000136 | -0.272 | *Col3a1* | 0.000008 | -0.1114 |
| *Cd14* | 0.000049 | 0.3031 | *Trem1* | 0.000182 | 0.219 | *Cxcl12* | 0.000008 | -0.0925 |
| *Cxcr2* | 0.000066 | 0.3311 | *Stat3* | 0.000184 | 0.0429 | *Col1a1* | 0.000009 | -0.1287 |
| *Trem1* | 0.000073 | 0.3327 | *Dcn* | 0.0002 | -0.1388 | *Cd68* | 0.00001 | 0.187 |
| *Wfdc21* | 0.00008 | 0.4754 | *Cd14* | 0.000209 | 0.1082 | *Tlr2* | 0.000015 | 0.1502 |
| *Cxcl3* | 0.000092 | 0.4199 | *Tim2* | 0.000278 | -0.1003 | *Fcgr1* | 0.000018 | 0.1277 |
| *Itgb5* | 0.000093 | -0.2112 | *Itgam* | 0.000309 | 0.1003 | *Hspg2* | 0.00002 | -0.1568 |
| *Nlpr3* | 0.000099 | 0.3079 | *Nlpr3* | 0.000325 | 0.167 | *Cd69* | 0.000021 | 0.2088 |
| *Cd69* | 0.00011 | 0.2712 | *Il1a* | 0.000327 | 0.2433 | *Tyrobp* | 0.000026 | 0.1303 |
| *Cxcl2* | 0.000123 | 0.358 | *Il36g* | 0.0004 | 0.3589 | *Ctsb* | 0.000027 | 0.09216 |
| *Jak2* | 0.000166 | 0.1312 | *Il4ra* | 0.00048 | 0.09126 | *Irf5* | 0.000034 | 0.1265 |
| *H2-Dma* | 0.000169 | -0.2365 | *Apoe* | 0.000524 | -0.1796 | *Cd14* | 0.000038 | 0.1511 |
| *Wnt-10b* | 0.000186 | -0.4969 | *Csf1* | 0.000614 | 0.1357 | *Trem2* | 0.000045 | 0.3059 |
| *Hif1a* | 0.000188 | 0.1598 | *Cx3r1* | 0.000703 | -0.2688 | *Itgb2* | 0.000048 | 0.1978 |
| *Fcgrt* | 0.0003 | -0.2941 | *Jak2* | 0.000839 | 0.06636 | *Tnf* | 0.000053 | 0.207 |
| *Il36g* | 0.000314 | 0.3203 | *Il1b* | 0.000872 | 0.1285 | *Dcn* | 0.000066 | -0.2512 |
| *Selplg* | 0.000324 | 0.2322 | *Nfkbiz* | 0.000929 | 0.06355 | *Col6a1* | 0.000083 | -0.2284 |
| *Il4ra* | 0.00037 | 0.1257 | *Tlr6* | 0.000932 | 0.1673 | *Tgfbr3* | 0.000088 | -0.1637 |
| *Mmp8* | 0.000389 | 0.2898 | *Eno1* | 0.000968 | 0.05611 | *C3ar* | 0.000093 | 0.1126 |
| *Il1a* | 0.000437 | 0.3307 | *Fap* | 0.001038 | -0.1421 | *Il1b* | 0.000094 | 0.1622 |
| *C5ar* | 0.000445 | 0.206 | *Il17ra* | 0.001052 | 0.0977 | *Mmp12* | 0.000138 | -0.1322 |
| *Apoe* | 0.000624 | -0.2117 | *Relb* | 0.001111 | 0.07728 | *Ctsz* | 0.000149 | 0.09127 |
| *Fgf7* | 0.000669 | -0.353 | *Selenop* | 0.001231 | -0.1269 | *Col5a1* | 0.000149 | -0.1722 |
| *Pigf* | 0.000776 | -0.1013 | *Tmem119* | 0.001413 | -0.1256 | *Fn1* | 0.000169 | -0.1506 |
| *Ccl4* | 0.000832 | 0.3715 | *Col5a1* | 0.001465 | -0.08864 | *Itgam* | 0.00017 | 0.1064 |
| *Tgfbr3* | 0.00097 | -0.1807 | *Bgn* | 0.001551 | -0.0689 | *Cxcr4* | 0.000209 | 0.1445 |
| *Ndufa1* | 0.001002 | -0.07342 | *Nfkb2* | 0.00166 | 0.09885 | *Timp3* | 0.000296 | -0.2874 |

| | | | | | | | | |
|---|---|---|---|---|---|---|---|---|
| *Cxcl12* | 0.001032 | -0.1381 | *Itgb5* | 0.001668 | -0.08094 | *Marco* | 0.000328 | 0.219 |
| *Il1b* | 0.001065 | 0.2524 | *Ccl4* | 0.001821 | 0.2002 | *Tgfb1* | 0.000384 | 0.1156 |
| *Nfkbiz* | 0.001132 | 0.1191 | *Mrc1* | 0.002099 | 0.1357 | *Mmp8* | 0.000388 | 0.3635 |
| *Dcn* | 0.001286 | -0.157 | *Mmp2* | 0.002231 | -0.145 | *Nr1h3* | 0.000473 | 0.2182 |
| *Mgl2* | 0.001437 | -0.3164 | *Col6a1* | 0.002376 | -0.09018 | *Tek* | 0.000473 | -0.3031 |
| *Ap2a2* | 0.001715 | -0.1254 | *Fcgrt* | 0.002425 | -0.1211 | *Itgb1* | 0.000493 | -0.06211 |
| *Fgf2* | 0.001749 | -0.2305 | *Itgb2* | 0.00252 | 0.07233 | *Fap* | 0.000569 | -0.1802 |
| *Stat2* | 0.001757 | -0.09352 | *Cxcl3* | 0.002651 | 0.3101 | *Mgl2* | 0.000721 | -0.2491 |
| *Mmp2* | 0.00184 | -0.1772 | *Vegfa* | 0.003208 | 0.0959 | *Gusb* | 0.000818 | 0.07138 |
| *Igf1* | 0.001945 | -0.3135 | *Angpt2* | 0.003322 | 0.1536 | *Plod* | 0.00084 | -0.09233 |
| *Nfkbia* | 0.002165 | 0.1513 | *Col1a2* | 0.003546 | -0.09114 | *C5ar* | 0.001026 | 0.1294 |
| *Tek* | 0.002351 | -0.2103 | *Cd86* | 0.003824 | 0.1458 | *Ctsl* | 0.001106 | 0.2317 |
| *Sell* | 0.002497 | 0.176 | *Nfkbia* | 0.004229 | 0.08654 | *Adgre1* | 0.001324 | 0.1019 |
| *Ndufc2* | 0.002501 | -0.1353 | *Cxcr2* | 0.004445 | 0.2448 | *Il17ra* | 0.001371 | 0.07895 |
| *Psme1* | 0.002594 | -0.08362 | *Jak1* | 0.004687 | 0.05138 | *Pdgfra* | 0.001435 | -0.08696 |
| *Tlr4* | 0.002825 | 0.1121 | *Tnf* | 0.005408 | 0.189 | *Wfdc17* | 0.001511 | 0.09883 |
| *Tyrobp* | 0.002974 | 0.207 | *Tgfb3* | 0.006256 | -0.1676 | *Nlpr3* | 0.00173 | 0.1804 |
| *S1pr4* | 0.003056 | -0.308 | *Hif1a* | 0.006398 | 0.06381 | *Psma4* | 0.001797 | 0.1345 |
| *Cd9* | 0.003057 | 0.1027 | *Rel* | 0.009637 | 0.08776 | *Nfkbia* | 0.001849 | 0.08747 |
| *Col5a1* | 0.003278 | -0.2613 | | | | *Cd52* | 0.001992 | 0.1207 |
| *S1pr2* | 0.003778 | -0.1337 | | | | *Cd80* | 0.002018 | 0.07633 |
| *Timp2* | 0.003884 | -0.1718 | | | | *Tgfb2* | 0.002236 | -0.1041 |
| *Mmp9* | 0.004007 | 0.2587 | | | | *Psme1* | 0.00226 | 0.054 |
| *Stat5b* | 0.004017 | -0.1028 | | | | *Tlr7* | 0.002301 | 0.1384 |
| *Asprv1* | 0.004155 | 0.2461 | | | | *Vcan* | 0.00318 | -0.13 |
| *Selenop* | 0.004408 | -0.2264 | | | | *Fcgrt* | 0.003322 | -0.0901 |
| *Wfdc17* | 0.004821 | 0.1546 | | | | *Hif1a* | 0.003554 | -0.06382 |
| *Klf4* | 0.004952 | -0.1267 | | | | *Ly69* | 0.004082 | 0.1976 |
| *Ctsb* | 0.005021 | -0.07897 | | | | *Stat3* | 0.004177 | -0.06242 |
| *Angpt1* | 0.005119 | -0.8333 | | | | *Tlr4* | 0.004219 | 0.0834 |
| *Cd80* | 0.005243 | 0.2139 | | | | *P2ry13* | 0.00435 | 0.09517 |

| | | | | | | | | |
|---|---|---|---|---|---|---|---|---|
| *Clta* | 0.005321 | -0.05317 | | | | *Ccr5* | 0.004529 | 0.06946 |
| *Itgb4* | 0.005428 | -0.9401 | | | | *Ly6c1* | 0.004537 | -0.1458 |
| *Cox5a* | 0.005573 | -0.083 | | | | *Stx5a* | 0.004629 | -0.06741 |
| *Irf4* | 0.005635 | -0.1845 | | | | *Pparg* | 0.005031 | 0.1703 |
| *Mrc1* | 0.00572 | -0.1238 | | | | *Clec7a* | 0.005194 | 0.1178 |
| *Cbr2* | 0.006169 | -0.3108 | | | | *Cd86* | 0.005258 | 0.09626 |
| *Spp1* | 0.006323 | 0.3553 | | | | *Itgal* | 0.005846 | 0.1598 |
| *Itgam* | 0.00643 | 0.1794 | | | | *Agr1a* | 0.005867 | -0.14 |
| *Cd163* | 0.00688 | -0.153 | | | | *Chil3* | 0.005992 | 0.3074 |
| *Tlr2* | 0.006884 | 0.2149 | | | | *Arg1* | 0.006103 | 0.3367 |
| *Pdgfrb* | 0.007536 | -0.128 | | | | *Vegfa* | 0.00613 | 0.1159 |
| *Fap* | 0.007697 | -0.2559 | | | | *Retnla* | 0.006681 | -0.6347 |
| *Axl* | 0.007777 | -0.1067 | | | | *Ccl4* | 0.006713 | 0.1713 |
| *Ly6c1* | 0.007919 | -0.137 | | | | *Irf7* | 0.006863 | 0.1254 |
| *Myd88* | 0.008542 | 0.0393 | | | | *Cxcl3* | 0.006923 | 0.4286 |
| *Uqcrq* | 0.008745 | -0.07996 | | | | *Psma3* | 0.007193 | 0.04564 |
| *Col3a1* | 0.009134 | -0.1988 | | | | *Tie1* | 0.007781 | -0.1717 |
| *Psme2* | 0.009259 | -0.06806 | | | | *Ndufa1* | 0.008039 | 0.07993 |
| *Gusb* | 0.00927 | -0.1125 | | | | *Col1a2* | 0.008861 | -0.04918 |
| *Csf1r* | 0.009289 | -0.1444 | | | | *Txnip* | 0.009226 | -0.1094 |
| *Ccl8* | 0.009532 | -0.1467 | | | | *Trem1* | 0.009507 | 0.3519 |

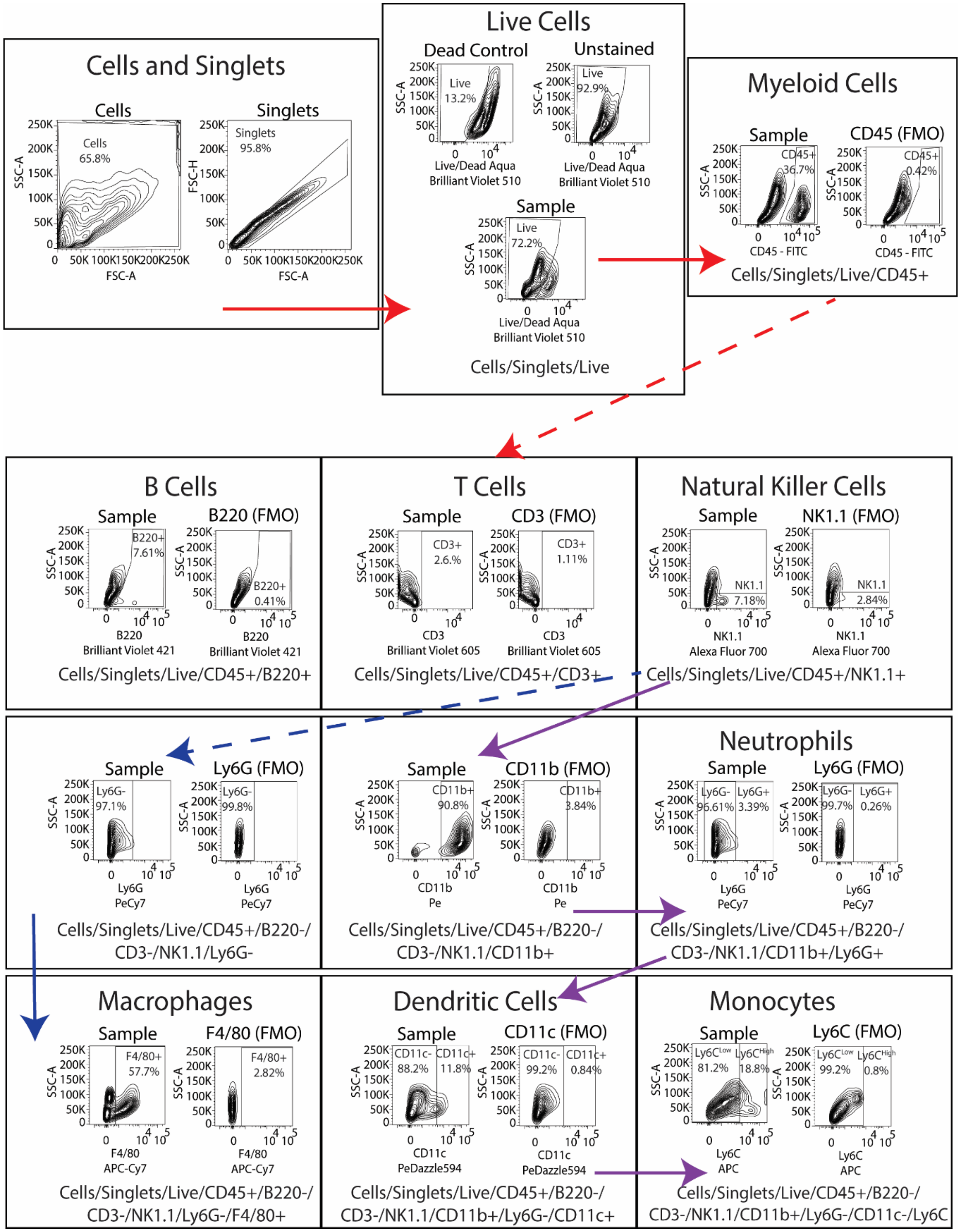

**Suppl. Figure 6.** Leukocyte presence gating strategy.

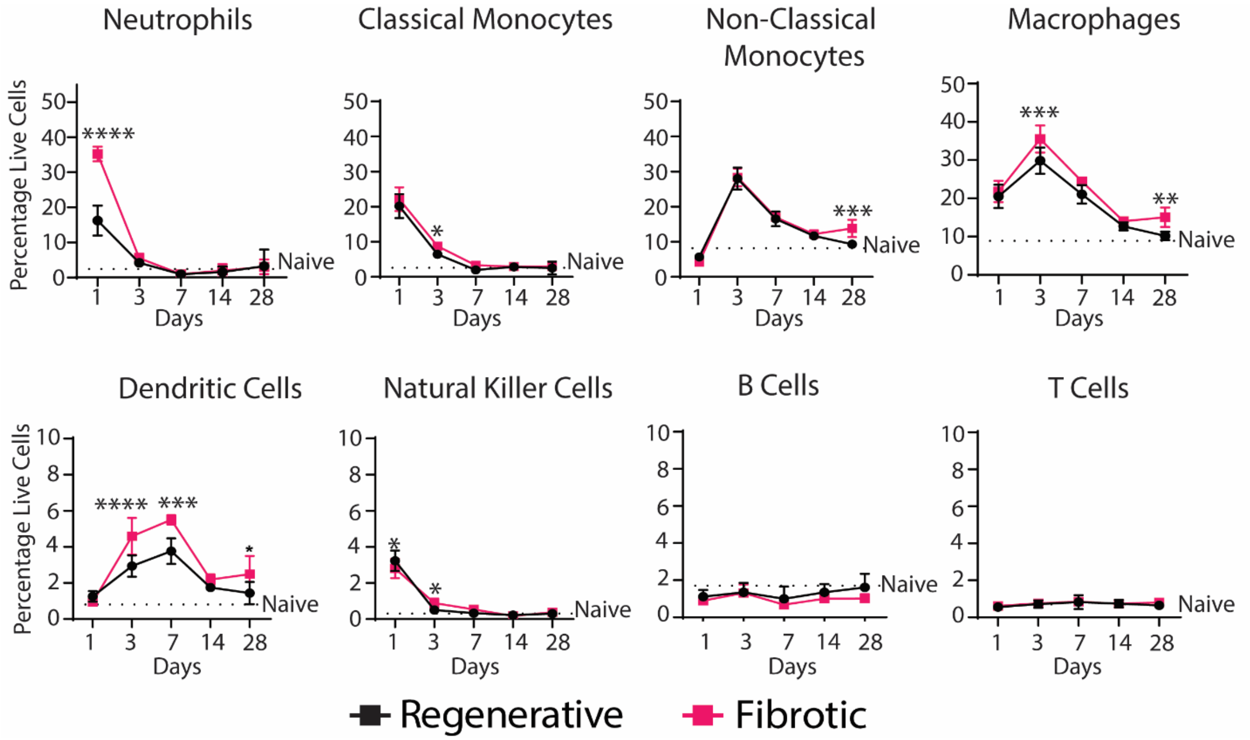

**Suppl. Figure 7.** Leukocyte accumulation in muscle over 28 days via flow cytometry. Two-Way ANOVA, Sidak's Post Hoc. N = 6, bars show mean ± SD p<0.0.5 * ; p < 0.01 ** ; p < 0.001 *** ; p < 0.0001 ****.

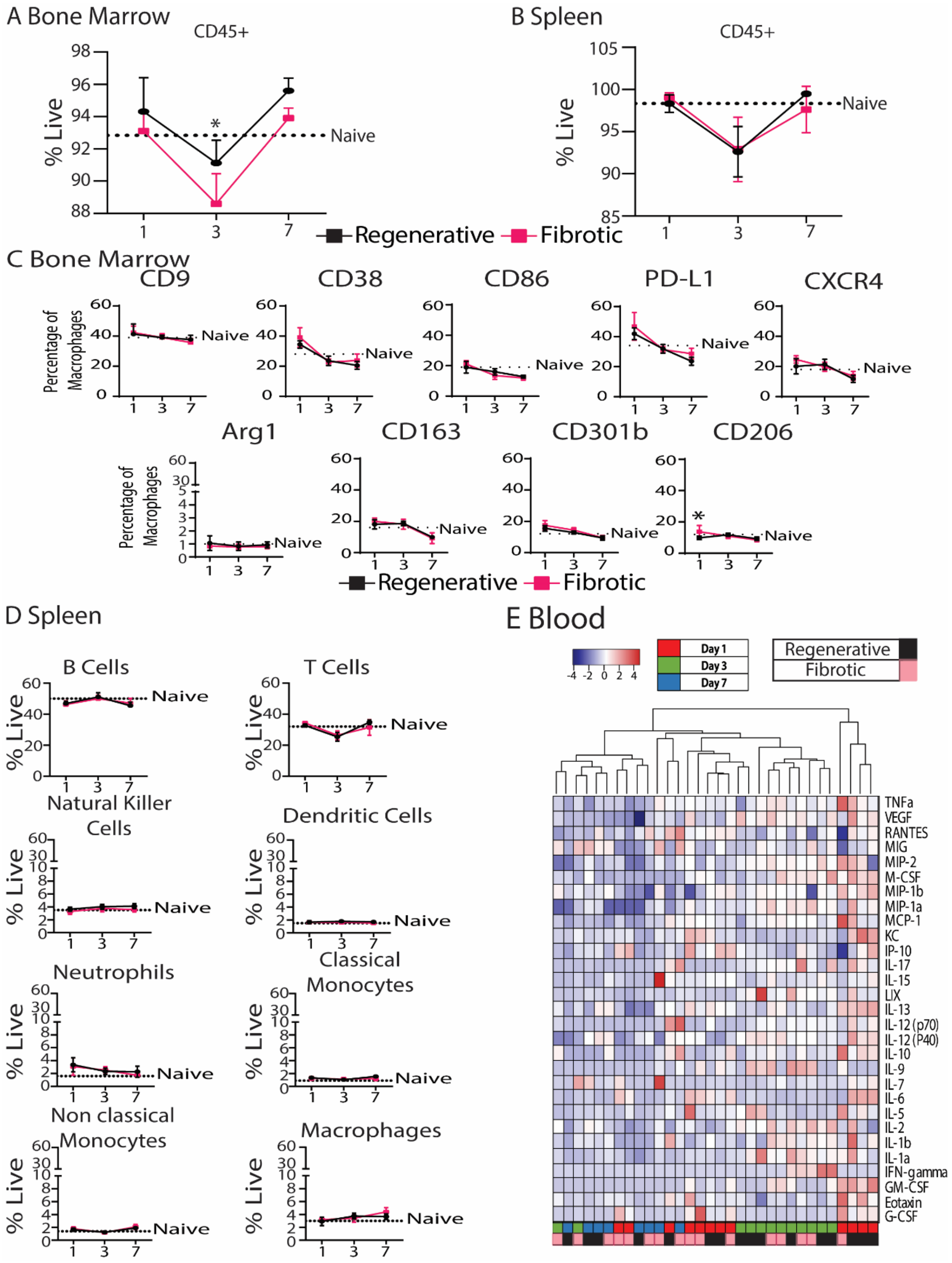

**Suppl. Figure 8. A)** CD45 expression in bone marrow. **B)** CD45 expression in spleen. **C)** Macrophage phenotype in bone marrow over 7 days. **D)** Immune cell presence in spleen over 7 days. Two-Way ANOVA, Sidak's Post Hoc. **E)** Analytes in blood serum. N = 6, bars show mean ± SD. p<0.0.5 * ; p < 0.01 ** ; p < 0.001 *** ; p < 0.0001 ****

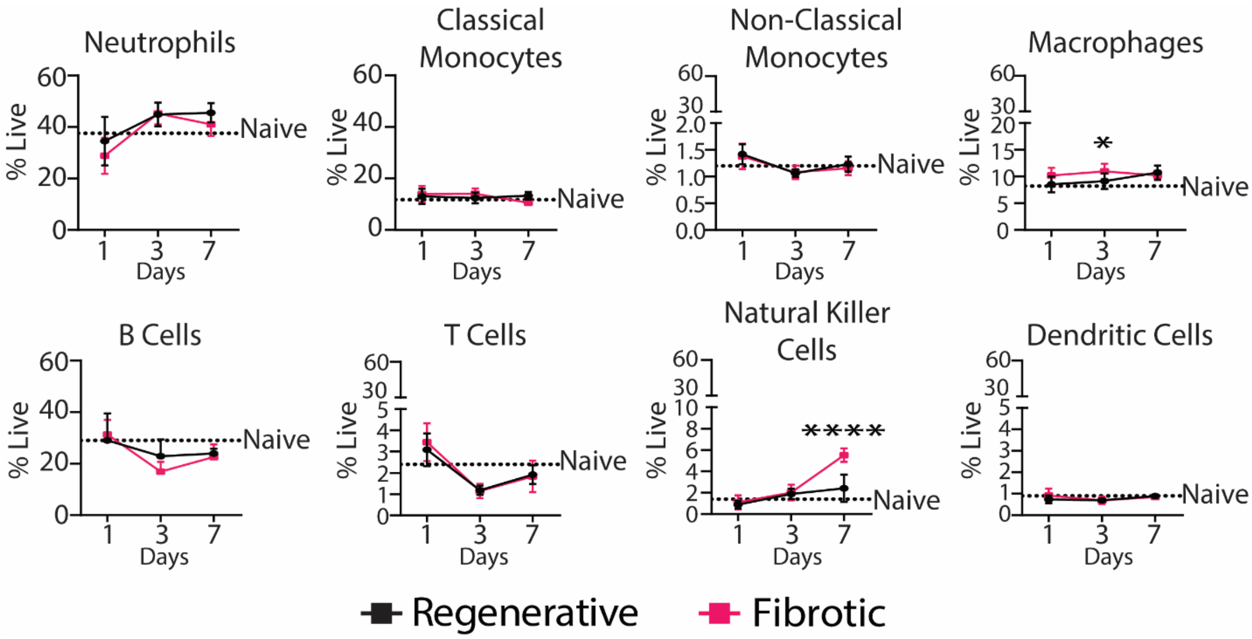

**Suppl. Figure 9.** Immune cell presence in the bone marrow over 28 days via flow cytometry. Two-Way ANOVA, Sidak's Post Hoc. N = 6, bars show mean ± SD. p<0.0.5 * ; p < 0.01 ** ; p < 0.001 *** ; p < 0.0001 ****

**Suppl. Table 4.** List of analytes in Luminex panel.

| | | | |
|---|---|---|---|
| G-CSF | IL-4 | LIF | MIP-1a |
| Eotaxin | IL-5 | IL-13 | MIP-1b |
| GM-CSF | IL-6 | LIX | M-CSF |
| IFNg | IL-7 | IL-15 | MIP-2 |
| IL-1a | IL-9 | IL-17 | MIG |
| L-1b | IL-10 | IP-10 | RANTES |
| IL-2 | IL-12 (p40) | KC | VEGF |
| IL-3 | IL-12 (p70) | MCP-1 | TNFa |

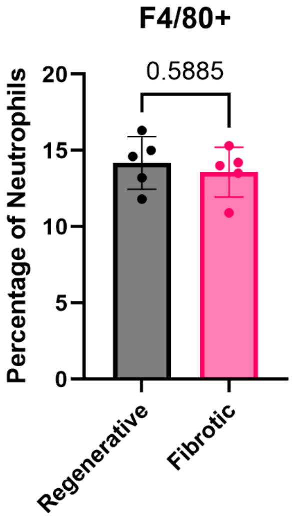

**Suppl. Figure 10.** F4/80 positive neutrophils (CD45+/BB20-/CD3-/NK1.1-/CD11b+/Ly6G+). T-test. N = 5, bars show mean ± SD p<0.0.5 * ; p < 0.01 ** ; p < 0.001 *** ; p < 0.0001 ****.

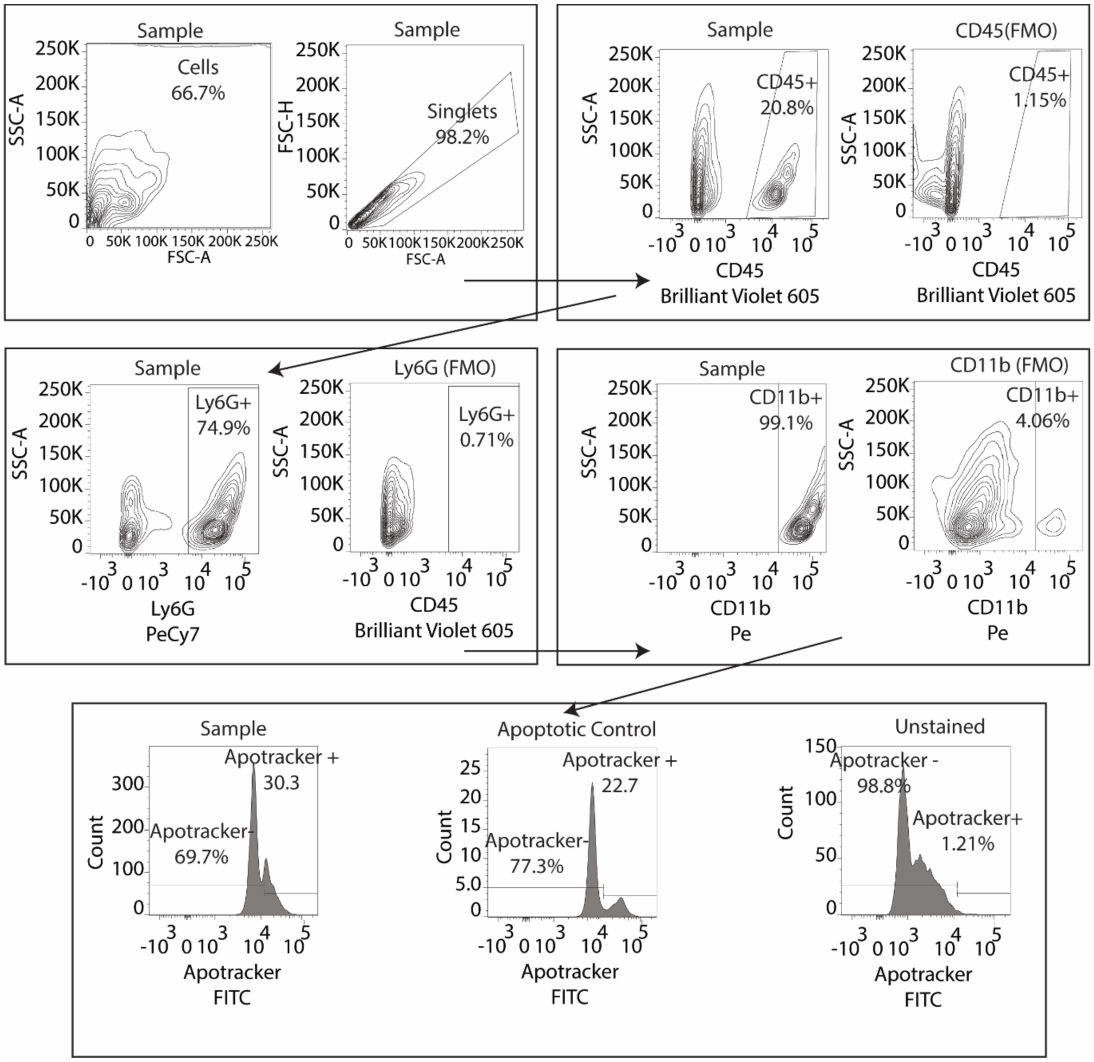

**Suppl. Figure 11.** Neutrophil apoptosis gating strategy.

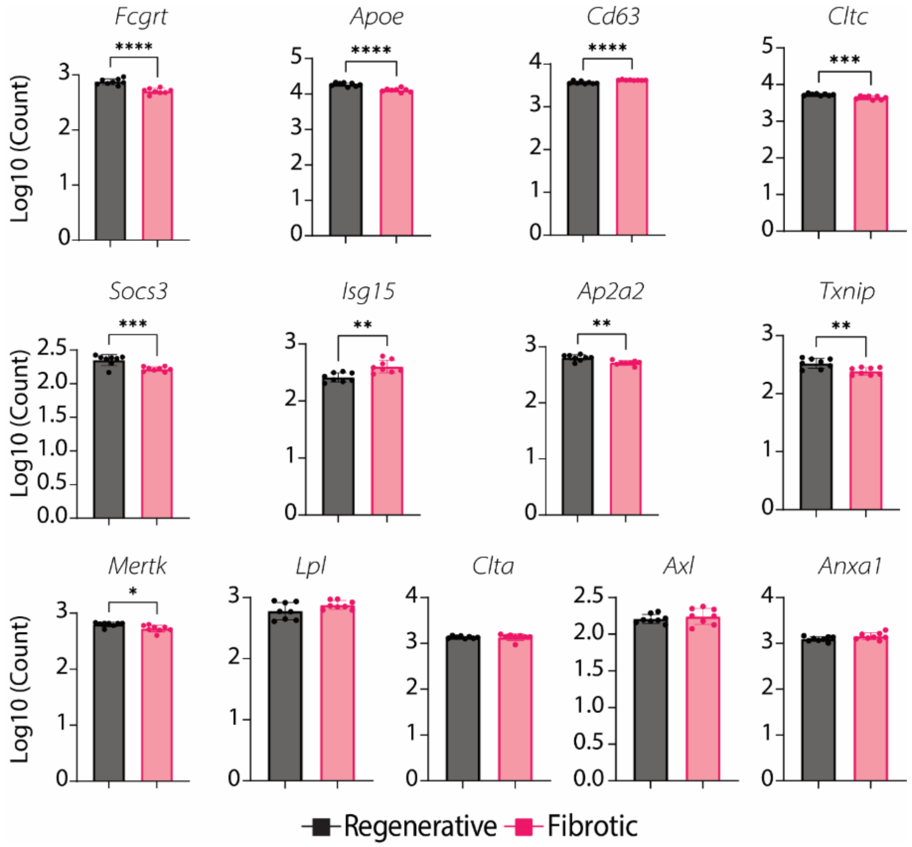

**Suppl. Figure 12.** Gene expression of phagocytosis related genes at day 3. T-test. N = 8, bars show mean ± SD p<0.0.5 * ; p < 0.01 ** ; p < 0.001 *** ; p < 0.0001 ****.

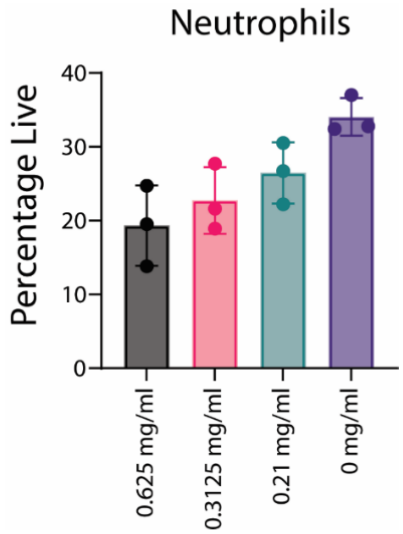

**Suppl. Figure 13.** Percentage of neutrophils in muscle following different dosages of tail-vein injected anti-Ly6G at day 1. One-Way ANOVA, Tukey's Post Hoc . N = 3, bars show mean ± SD p<0.0.5 * ; p < 0.01 ** ; p < 0.001 *** ; p < 0.0001 ****.

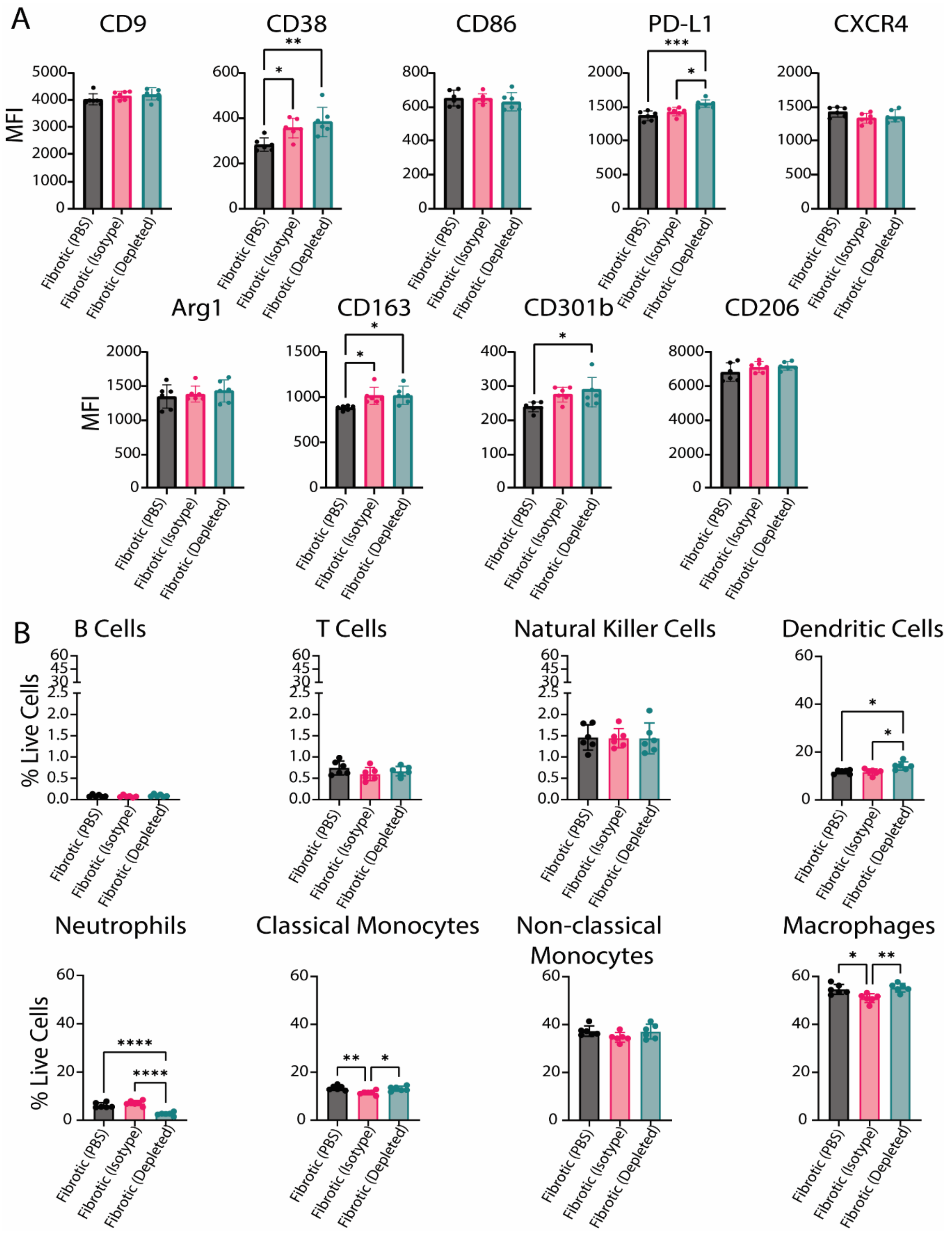

**Suppl. Figure 14. A)** Macrophage phenotype in muscle on day 3 follow neutrophil depletion compared to isotype control and PBS control. **B)** Immune cell presence in muscle on day 3. One-Way ANOVA, Tukey's Post Hoc. N = 6, bars show mean ± SD p<0.0.5 * ; p < 0.01 ** ; p < 0.001 *** ; p < 0.0001 ****

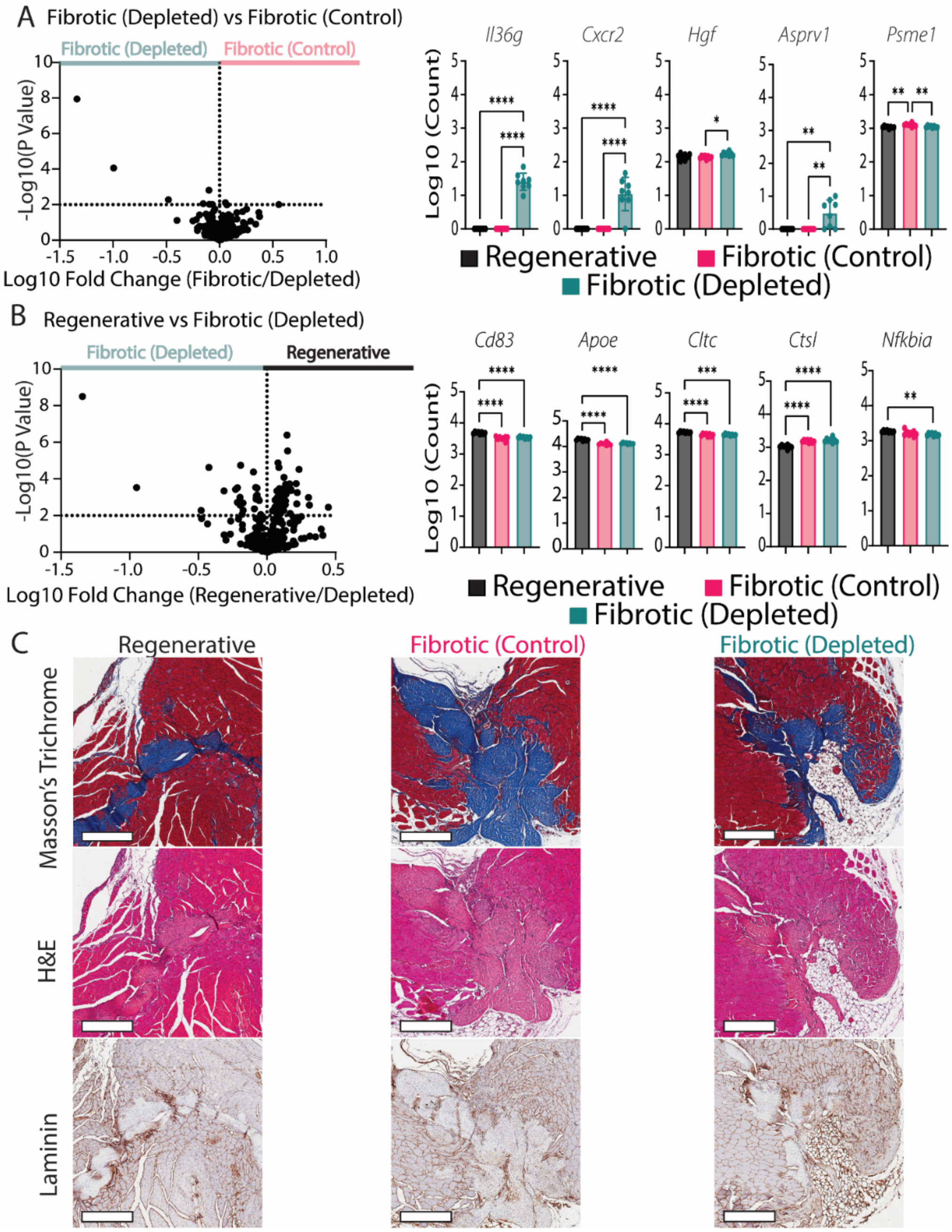

**Suppl. Figure 15. A)** Volcano plot showing fold change in gene expression between the fibrotic (neutrophil-depleted) versus fibrotic (control) groups (multiple t-tests, p<0.01 cutoff) and the top 5 DEGs (one-way ANOVA, Tukey's post hoc) of FACS-sorted macrophages at day 3. **B)** Volcano plot of fibrotic (neutrophil-depleted) versus regenerative groups (multiple t-tests, p<0.01 cutoff) and top 5 DEGs (one-way ANOVA, Tukey's post hoc) of FACS-sorted macrophages at day 7. **C)** Representative images of Masson's trichrome, H&E and laminin stains (scale bar 200 um). N = 8, bars show mean ± SD * p<0.0.5; ** p < 0.01; *** p < 0.001; p < 0.0001 ****.

**Suppl. Table. 5.** List of differentially expressed genes (DEGs), p-value and fold change (Fibrotic vs Regenerative).

| Regenerative vs Fibrotic | | | Depleted vs Fibrotic | | | Depleted vs Regenerative | | |
|---|---|---|---|---|---|---|---|---|
| **Gene Name** | **P Value** | **Fold Change** | **Gene Name** | **P Value** | **Fold Change** | **Gene Name** | **P Value** | **Fold Change** |
| *Mgl2* | 0.000001 | -0.3961 | *Il6* | <0.000001 | -1.339 | *Il36g* | <0.000001 | -1.344 |
| *Ccr2* | 0.000003 | -0.1704 | *Cxcr2* | 0.000086 | -0.994 | *Cd83* | <0.000001 | 0.1461 |
| *Cx3cr1* | 0.000005 | -0.2419 | *Hgf* | 0.001545 | -0.09987 | *Apoe* | 0.000003 | 0.1506 |
| *Ctsl* | 0.000005 | 0.157 | *Asprv1* | 0.005251 | -0.4794 | *Cltc* | 0.000013 | 0.0803 |
| *Retnla* | 0.000005 | -0.6347 | *Psme1* | 0.007779 | 0.05541 | *Ctsl* | 0.000018 | -0.1919 |
| *Mrc1* | 0.000011 | -0.1637 | *Csf1* | 0.0089 | -0.154 | *Nfkbia* | 0.000023 | 0.08708 |
| *Cbr2* | 0.000012 | -0.1924 | *Lpl* | 0.009568 | -0.09513 | *Csf1* | 0.000023 | -0.4229 |
| *Cd83* | 0.000013 | -0.171 | *Ccr2* | 0.009721 | -0.06697 | *Fcgrt* | 0.00003 | 0.2342 |
| *Fcgrt* | 0.000014 | -0.1686 | *Acta2* | 0.009873 | 0.556 | *Tyrobp* | 0.000042 | -0.09877 |
| *Apoe* | 0.000018 | -0.1613 | | | | *Nfkbiz* | 0.000082 | 0.1243 |
| *Cd68* | 0.000029 | 0.09993 | | | | *Cbr2* | 0.000179 | 0.2133 |
| *Tlr2* | 0.000076 | -0.1112 | | | | *Cx3cr1* | 0.000186 | 0.1646 |
| *Spp1* | 0.000077 | 0.3169 | | | | *Mrc1* | 0.000199 | 0.1358 |
| *Cd63* | 0.000096 | 0.06085 | | | | *Psma3* | 0.000292 | -0.08138 |
| *Nfkbiz* | 0.000133 | -0.1319 | | | | *Rela* | 0.000293 | 0.1354 |
| *Nr1h3* | 0.000146 | 0.2901 | | | | *Cxcr2* | 0.000294 | -0.9494 |
| *Cltc* | 0.000296 | -0.08637 | | | | *Sell* | 0.000301 | -0.2147 |
| *Pparg* | 0.000393 | 0.2072 | | | | *Uqcrq* | 0.00031 | -0.07279 |
| *Tyrobp* | 0.000423 | 0.1178 | | | | *H2-Dma* | 0.00034 | 0.0904 |
| *Cxcr4* | 0.000464 | -0.1013 | | | | *Ap2a2* | 0.000344 | 0.1272 |
| *H2-Ab1* | 0.000486 | -0.13 | | | | *Csf1r* | 0.000351 | 0.1019 |
| *Cd74* | 0.000623 | -0.1257 | | | | *Chil3* | 0.00037 | -0.2288 |
| *Psma4* | 0.000725 | 0.08692 | | | | *Fn1* | 0.000377 | 0.1746 |
| *Socs3* | 0.000764 | -0.1338 | | | | *H2-Eb1* | 0.000429 | 0.176 |

| | | | | | | | | |
|---|---|---|---|---|---|---|---|---|
| *Tmem119* | 0.000789 | -0.218 | | | | *Stat3* | 0.000451 | 0.07321 |
| *Uqcrq* | 0.000803 | 0.09646 | | | | *Spp1* | 0.000453 | -0.3123 |
| *Csf1* | 0.000887 | 0.269 | | | | *Tlr5* | 0.00056 | 0.2211 |
| *Nfkb1* | 0.000892 | -0.07894 | | | | *Ccr2* | 0.000595 | 0.1035 |
| *Itgb1* | 0.001129 | -0.0622 | | | | *Tlr7* | 0.000671 | 0.1397 |
| *Tlr9* | 0.001291 | -0.1475 | | | | *Cd68* | 0.000762 | -0.0718 |
| *C5ar* | 0.0013 | -0.1053 | | | | *Psma4* | 0.000847 | -0.07733 |
| *Tlr5* | 0.001406 | -0.172 | | | | *Il4ra* | 0.000934 | 0.1017 |
| *Ctsz* | 0.001416 | 0.05797 | | | | *Adgre1* | 0.001009 | 0.07577 |
| *Itga4* | 0.001428 | -0.08534 | | | | *Cd9* | 0.00101 | -0.1959 |
| *H2-Aa* | 0.001606 | -0.09895 | | | | *Mgl2* | 0.001025 | 0.3029 |
| *Isg15* | 0.001657 | 0.1891 | | | | *Relb* | 0.001124 | 0.08095 |
| *Ap2a2* | 0.001876 | -0.09046 | | | | *Ndufa1* | 0.001186 | -0.07392 |
| *Fn1* | 0.002097 | -0.2083 | | | | *Ctsz* | 0.001189 | -0.06474 |
| *Psme1* | 0.002171 | 0.06873 | | | | *Hif1a* | 0.001198 | 0.1235 |
| *H2-Dma* | 0.002336 | -0.07428 | | | | *Ctsd* | 0.001255 | -0.07972 |
| *H2-DMb1* | 0.002436 | -0.1103 | | | | *Psme2* | 0.001404 | -0.0596 |
| *Jak1* | 0.002665 | -0.06437 | | | | *Itga4* | 0.001423 | 0.05935 |
| *Stat1* | 0.002685 | 0.0838 | | | | *Cd63* | 0.001512 | -0.05567 |
| *Rel* | 0.002926 | -0.1132 | | | | *Socs3* | 0.001561 | 0.1348 |
| *Txnip* | 0.002951 | -0.136 | | | | *C5ar* | 0.001823 | 0.09383 |
| *Ndufa1* | 0.003166 | 0.08786 | | | | *Nr1h3* | 0.001901 | -0.215 |
| *Dcn* | 0.003884 | -0.1885 | | | | *Pparg* | 0.002042 | -0.1754 |
| *Cox5a* | 0.004334 | 0.05389 | | | | *Cxcr4* | 0.002209 | 0.08739 |
| *P2ry12* | 0.004372 | -0.1337 | | | | *Tmem119* | 0.00231 | 0.2176 |

| | | | | | | | | |
|---|---|---|---|---|---|---|---|---|
| *Mki67* | 0.004374 | -0.1856 | | | | *Cd74* | 0.002542 | 0.1339 |
| *Fcgr1* | 0.004778 | 0.1066 | | | | *Tlr9* | 0.002551 | 0.1074 |
| *Eno1* | 0.004801 | 0.0785<br>5 | | | | *Lpl* | 0.002663 | -0.1948 |
| *Irf4* | 0.005041 | -0.1479 | | | | *S1pr1* | 0.002665 | 0.3096 |
| *Clec10a* | 0.005349 | -0.1447 | | | | *Jak1* | 0.003454 | 0.0588<br>3 |
| *Csf1r* | 0.005696 | -<br>0.0966<br>3 | | | | *Retnla* | 0.003517 | 0.4469 |
| *Tlr3* | 0.005723 | -0.1791 | | | | *Clec7a* | 0.00425 | -<br>0.0917<br>1 |
| *Cd163* | 0.005798 | -0.2508 | | | | *Itgam* | 0.004433 | 0.0920<br>2 |
| *Klf4* | 0.006507 | -0.1046 | | | | *Cd6* | 0.005164 | -0.1742 |
| *Marco* | 0.007221 | 0.0746<br>4 | | | | *H2-Ab1* | 0.005229 | 0.1222 |
| *Adgre1* | 0.007511 | -0.1239 | | | | *Asprv1* | 0.005251 | -0.4794 |
| *Cd14* | 0.008578 | -<br>0.0450<br>1 | | | | *P2ry12* | 0.005305 | 0.1018 |
| *Pdgfb* | 0.009125 | -0.1774 | | | | *Itgb1* | 0.005547 | 0.041 |

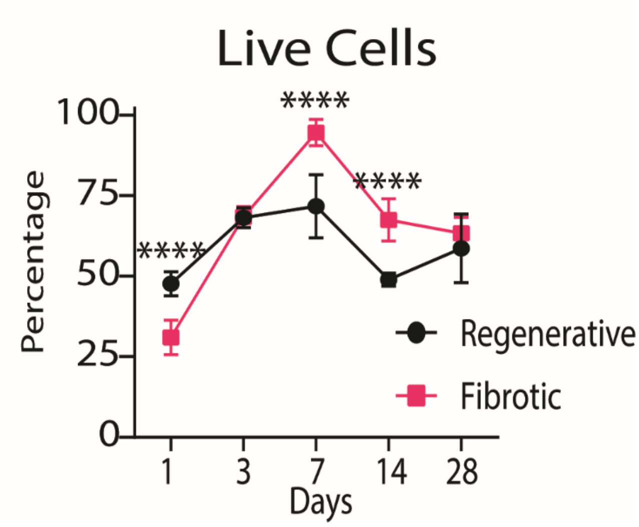

**Suppl. Figure 16.** Live cells in muscle over 28 days. Two-Way ANOVA, Sidak's Post Hoc. N = 6, bars show mean ± SD. p<0.0.5 * ; p < 0.01 ** ; p < 0.001 *** ; p < 0.0001 ****

### Suppl. Methods: MATLAB code for collagen staining quantification from Masson's trichrome images.

```
clc
clear
close all
File = imread("4 mm 1.tif");
figure;
imshow(File);
title('Original File');
Red = File(:,:,1); % Extracting Red Channel
Green = File(:,:,2); % Extracting Green Channel
Blue = File(:,:,3); % Extracting Blue Channel
JustBlue = Blue-Red/2-Green/2; % Removing background of other channels on blue
channel (aka collagen)
MeanJustBlue = mean(mean(JustBlue));
JustBlueMinusBackground = JustBlue < 2*MeanJustBlue; % Fine tuning blue
channel (scale factor of 2 manually adjusted)
figure;
imshow(JustBlueMinusBackground);
title('Collagen')
Overlay = imfuse(File, JustBlueMinusBackground);
figure;
subplot(2,2,1)
imshow(File)
subplot(2,2,2)
imshow(JustBlueMinusBackground)
subplot(2,2,[3,4])
imshow(Overlay)
BWBlue = imbinarize(Blue); % Represent all stained pixels, including NOT
collagen
BWBlue(BWBlue == 1) = [];
JustBlueMinusBackground(JustBlueMinusBackground == 1) = []; % Extract only
blue pixels (aka blue)
SizeBlue = length(JustBlueMinusBackground); % Calculates number of stained
pixels
SizeBWBlue = length(BWBlue); % Calculate number of blue pixels
Area = (SizeBlue/SizeBWBlue)*100; % Percentage of blue pixels in all stained
pixels
```
